# The minimizer Jaccard estimator is biased and inconsistent*

**DOI:** 10.1101/2022.01.14.476226

**Authors:** Mahdi Belbasi, Antonio Blanca, Robert S. Harris, David Koslicki, Paul Medvedev

## Abstract

**Motivation:** Sketching is now widely used in bioinformatics to reduce data size and increase data processing speed. Sketching approaches entice with improved scalability but also carry the danger of decreased accuracy and added bias. In this paper, we investigate the minimizer sketch and its use to estimate the Jaccard similarity between two sequences.

**Results:** We show that the minimizer Jaccard estimator is *biased* and *inconsistent*, which means that the expected difference (i.e., the bias) between the estimator and the true value is not zero, even in the limit as the lengths of the sequences grow. We derive an analytical formula for the bias as a function of how the shared *k*-mers are laid out along the sequences. We show both theoretically and empirically that there are families of sequences where the bias can be substantial (e.g. the true Jaccard can be more than double the estimate). Finally, we demonstrate that this bias affects the accuracy of the widely used mashmap read mapping tool.

**Availability:** Scripts to reproduce our experiments are available on GitHub [26].

## 1 Introduction

Sketching is a powerful technique to drastically reduce data size and increase data processing speed. Sketching techniques create a smaller representation of the full dataset, called a *sketch*, in a way that makes algorithms more efficient, ideally without much loss of accuracy. This property has led to sketching methods being increasingly used to meet the scalability challenges of modern bioinformatics datasets, though sometimes without understanding the detrimental effects on accuracy.

A thorough treatment of sketching in bioinformatics can be found in the excellent surveys of [29, 21], but we mention a few notable examples next. The seminal Mash paper [25] showed how estimating the Jaccard similarity of two sequences from their minhash sketches [3] enables clustering of sequence databases at unprecedented scale. The hyperloglog sketch [10] is used to compute genomic distances [1]; the modulo sketch [34] is used to search sequence databases [27]; strobemers [30] and minhash with optimal densification [36, 39] are used for sequence comparison; order minhash is used to estimate edit distance [19]; and count minsketch [5] is used for *k*-mer counting [6].

One of the most widely used sketches, which forms the basis of our work, is the minimizer sketch [34, 28], which selects, for each window of *w* consecutive *k*-mers, the *k*-mer with the smallest hash value. Minimizer sketches are used for transcriptome clustering [31] and error correction [32], as well as for seed generation by the Peregrine genome assembler [4] and the widely used minimap [16, 17] and mashmap [12, 13] aligners.

Just as with other sketching techniques, in order for the minimizer sketch to be useful, it must come with theoretical (or at least empirical) bounds on the loss of accuracy that results from its use. For instance, the minhash Jaccard estimator used by Mash has the property of being *unbiased* [3], i.e. its expected value is equal to the true Jaccard. Such a theoretical guarantee, however, cannot be assumed for other sketches. Here, we will consider the example of the *minimizer Jaccard estimate* [12, 13, 15], which computes the Jaccard similarity using minimizer sketches and forms the basis of the widely used mashmap [12, 13] aligner. This estimator is useful for sequence alignment because the minimizer sketch has the nice property that, roughly speaking, the sketch of a long string contains the sketches of all its substrings. However, its theoretical accuracy has not been studied and empirical evaluations have been limited.

In this paper, we study the accuracy of the minimizer Jaccard estimator *Ĵ*, both theoretically and empirically. We prove that *Ĵ* is in fact biased and inconsistent (i.e. the bias is not zero, and it remains so even as the sequences lengths grow). We derive an approximate formula for the bias that is accurate up to a vanishingly small additive error term, and give families of sequence pairs for which *Ĵ* is expected to be only between 40% to 63% of the true Jaccard. We then empirically evaluate the extent of the bias and find that in some cases, when the true Jaccard similarity is 0.90, the estimator is only 0.44. We also study both theoretically and empirically the bias of *Ĵ* for pairs of sequences generated by a simple mutation process and find that, while not as drastic, the bias remains substantial. Finally, we show that the bias affects the mashmap aligner by causing it to output incorrect sequence divergence estimates, with up to a 14% error. Our results serve as a cautionary tale on the necessity of understanding the theoretical and empirical properties of sketching techniques.

## 2 The minimizer sketch and minimizer Jaccard estimator

In this section, we will define the minimizer sketch [34, 28] and the Jaccard estimator derived from it [12]. Let *k* > 2 and *w* > 2 be two integers. This paper will assume that we are given two duplicate-free sequences *A* and *B* of *L k*-mers, with *L*⩾7(*w*+1). A sequence is *duplicate-free* if it has no duplicate *k*-mers, but *A* and *B* are allowed to share *k*-mers. These requirements on the sequences do not limit the general scope of our results. In particular, since we will show the existence of bias for these constrained cases, it immediately implies the existence of bias within broader families of sequences.

Let *A*_*i*_ denote the *k*-mer starting at position *i* of *A*, with *A*_0_ and *A*_*L*−1_ being the first and last *k*-mers, respectively. Let Sp^*k*^ (*A*) be the set of all *k*-mers in *A*. We define *I*(*A, B*) to be the number of *k*-mers shared between *A* and *B*, and *U* (*A, B*) to be the number of *k*-mers appearing in either *A* or *B*. Formally,

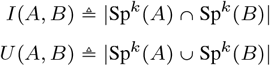

The Jaccard similarity between the sequences *A* and *B* is defined as

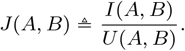

Suppose we have a hash function *h* that takes an element from the set of all *k*-mers and maps it to a real number drawn uniformly at random from the unit interval [0, 1]. Under this hash function, the probability of a collision is 0. We denote by *a*_*i*_ the hash value assigned to *k*-mer *A*_*i*_ and for integer *w* ⩾ 2 define the minimizer sketch of *A* as

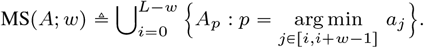

An element in MS (*A*; *w*) is called a *minimizer* of *A*. The minimizer intersection and the minimizer union of *A* and *B* are defined, respectively, as

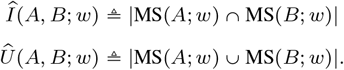

The minimizer Jaccard estimator between *A* and *B* is defined as

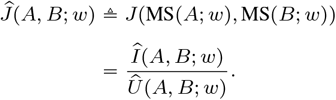

## 3 Main theoretical results

In this section, we state our main theoretical results and give some intuition behind them. We can think of the relationship between the shared *k*-mers of *A* and *B* as the subset of (*A*_0_, …, *A*_*L*−1_) × (*B*_0_, …, *B*_*L*−1_) that corresponds to pairs of equal elements; i.e., to pairs (*A*_*i*_, *B*_*j*_) with *A*_*i*_ *= B*_*j*_. Because *A* and *B* are duplicate-free, this relationship is a matching. We call this the *k-mer-matching* between *A* and *B*. Our main result is stated in terms of a term denoted by ℬ (*A, B*; *w*), which is a deterministic function of the window size *w* and of the *k*-mer-matching between *A* and *B*. We postpone the exact definition of ℬ (*A, B*; *w*) until Appendix A.1, since it requires the introduction of cumbersome notation. The main technical result of this paper is:

### Theorem 1.

*Let w* ⩾2, *k* ⩾ 2, *and L* ⩾ 7(*w*+1) *be integers. Let A and B be two duplicate-free sequences, each consisting of L k-mers. Then there exists* 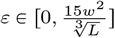 *such that*

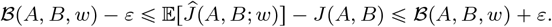

In other words, the difference between the expected value of the minimizer Jaccard estimator and the true Jaccard is ℬ(*A, B*; *w*), up to a vanishingly small additive error. We now investigate the value of the term *ℬ*, which approximates the bias. First, we can show that for padded sequences, ℬ(*A, B*; *w*) < 0, except that ℬ(*A, B*; *w*) = 0 when *J* (*A, B*) = 0. We say two sequences are padded if they do not share any minimizers in the first or l ast *w k* -mers. (We note that the effect of padding becomes negligible for longer sequences.)

### Theorem 2.

*Let w* ⩾ 2, *k* ⩾ 2, *and L* ⩾ 7(*w*+1) *be integers. Let A and B be two duplicate-free padded sequences, each consisting of L k-mers. Then ℬ* (*A, B*; *w*) < 0 *unless J* (*A, B*) = 0; *when J* (*A, B*) = 0, *we have ℬ*(*A, B*; *w*) = 0.

Moving forward, we may omit *A, B*, and *w* from our notation when they are obvious from the context. Theorems 1 and 2 state that *Ĵ* is biased for padded sequences as long as *ε* is sufficiently small (e.g. *L* is sufficiently large or *w* is sufficiently small). Here, we use “biased” in the statistical sense that 𝔼[*Ĵ*] ≠ *J*. Intuitively, *Ĵ* is biased because it depends on the layout of the shared *k*-mers along the sequences (i.e. on the *k*-mer-matching), while *J* only depends on the number of shared *k*-mers but not on their layout. Note that our results hold for any duplicate-free choice of *A* and *B* and do not assume any background distribution, e.g. that *A* is generated uniformly at random.

We illustrate the point with Examples 2a and 2b in Figure 1. In both examples, the expected size of *Î* is the probability that *x* is a minimizer in *A* and in *B* plus the probability that *y* is a minimizer in *A* and in *B*. These two probabilities are equal to each other in these examples and 𝔼[*Î*] = 2*τ*, for some *τ*. When *w* = 2, in Example 2a, *τ* is the probability that *a*_1_ is the smallest hash value out of five independently chosen hash values *a*_0_, *a*_1_, *a*_2_, *b*_0_, and *b*_2_. In Example 2b, however, *b*_2_ = *a*_2_, and *τ* is the probability that *a*_1_ is the smallest hash value out of four independently chosen hash values (*a*_0_, *a*_1_, *a*_2_, *b*_0_). Hence, the values of *τ* are different in the two examples, and therefore 𝔼[*Î*] is also different.

**Fig. 1:**
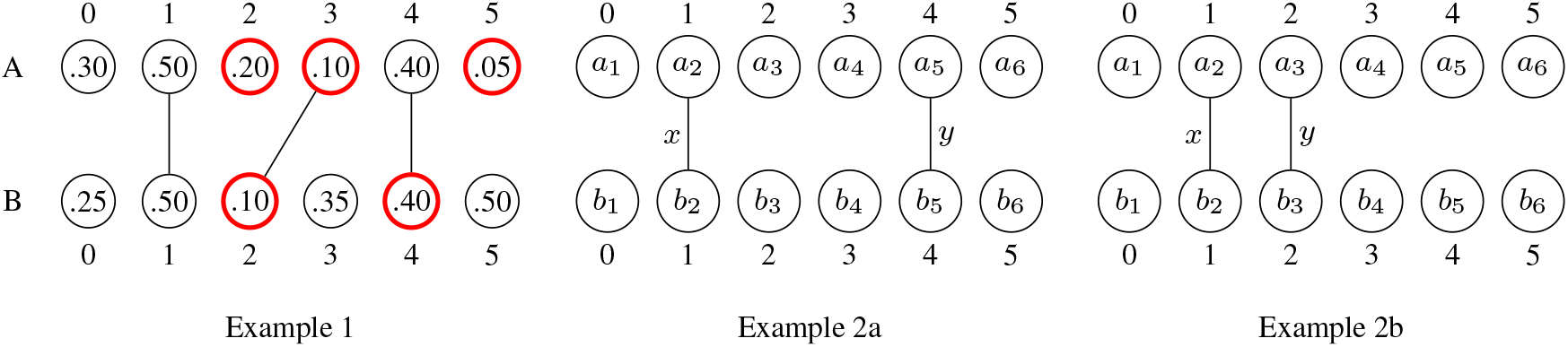
Examples of the Jaccard and the minimizer Jaccard estimator. Each example shows the *k*-mers of a sequence *A* on top, the *k*-mers of a sequence *B* on the bottom, and lines connecting *k*-mers show the *k-mer-matching* between *A* and *B*. Each *k*-mer is labeled by its hash value. In Example 1, *J* (*A, B*) = 1/3. The minimizers for *w* = 3 are circled in bold red. Here, *Î (A, B;* 3) = 1, *Û* (*A, B;* 3) = 4, and *Ĵ (A, B;* 3) = 1/4. Examples 2a and 2b give intuition for why the minimizer Jaccard estimator is biased. Here, *a*_*i*_ refers to the hash value assigned to position *i* and *x* and *y* are *k*-mers shared between *A* and *B*. The expected minimizer Jaccard for *w* = 2 is different in the two examples but the Jaccard is not (*J* = 0.2); hence the expected minimizer Jaccard cannot be equal to the true Jaccard.

The discrepancy on 𝔼[*Î*] turns out to be crucial since it induces a bias. Specifically, as part of the proof of Theorem 1, we will show that 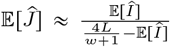, and, since the difference between the expected sizes of the minimizer intersections varies for the two examples, we have that 𝔼[*Ĵ*] is also different; in particular, 𝔼[*Ĵ*] is affected by the layout of the *k*-mer-matching. Note, however, that the Jaccard similarity in both examples is the same, with *J* = 0.2, leading to the intuition that *Ĵ* is biased when *w* = 2. Theorems 1 and 2 show that this bias extends beyond this contrived example and holds for most sequences of interest.

Next, we consider the value of ℬ(*A, B*; *w*) for some more concrete families of sequence pairs. First, consider the case where any pair of *k*-mers that are shared between *A* and *B* are separated by at least *w* positions. This may approximately happen in practice when *A* and *B* are biologically unrelated and the *k*-mer matches are spurious. Formally, we say two padded sequences *A* and *B* are *sparsely-matched* if for all *p* and *q* such that *A*_*p*_ = *B*_*q*_, {*A*_*p*−*w*_, …, *A*_*p*−1_, *A*_*p*−1_, …, *A*_*p*+*w*_} ∉ Sp^*k*^ (*B*), and {*B*_*q*−*w*_, …, *B*_*q*−1_, *B*_*q*−1_, …, *B*_*q*−*w*_} ∉ Sp^*k*^ (*A*). In such a case, one could imagine that since the shared *k*-mers do not interfere with each other’s windows, the estimator might be unbiased. It turns out this is not the case.

### Theorem 3.

*Let w* ⩾ 2, *k* ⩾ 2, *and L* ⩾ 7(*w*+1) *be integers. Let A and B be two duplicate-free, padded, sparsely-matched sequences, each consisting of L k-mers. Then* 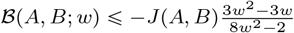.

A direct consequence of combining this with Theorem 1 is that for sparsely-matched sequences with *J* (*A, B*) > 0,

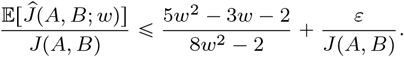

For example, for *w* = 20 and sufficiently long sequence pairs with a fixed (i.e. independent of *L* or *w*) Jaccard similarity, *Ĵ* is at most 61% of the true Jaccard. The bias cannot be fixed by changing *w*, since at *w* = 2, *Ĵ* is at most 40% of *Ĵ*, and, as *w* grows, *Ĵ* is at most 63% of the true Jaccard. This example also shows that *Ĵ* is not only biased but also *inconsistent*, i.e. 𝔼[*Ĵ*] does not converge to *J* even as the sequences grow long.

Let us now consider the opposite side of the spectrum, where instead of being sparsely-matched, *A* and *B* are related by the simple mutation model (i.e. every position is mutated with some constant probability [2]). Deriving the bias for this case proved challenging, since the mutation process adds another layer of randomness. Instead, we derive the bias in a simpler deterministic version of this process, where there is a mutation every *g* positions, for some *g* > *w* +2*k*.

### Theorem 4.

*Let* 2 ⩽*w* < *k, g* > *w* + 2*k, and L* = 𝓁*g* + *k for some integer* 𝓁 ⩾ 1. *Let A and B be two duplicate-free sequences with L k-mers such that A and B are identical except that the nucleotides at positions k* − 1 + *ig, for i* = 0, …, 𝓁, *are mutated. Then*,

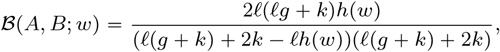

*where* 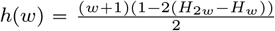 *and* 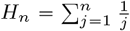 *denotes the n-th Harmonic number*.

We can use this theorem in combination with Theorem 1 to obtain a precise approximation of the bias of *Ĵ* for this family of sequences. For instance, taking *k* = 15, *w* = 10, *L* = 9992, and *g* = 43 yields that *Ĵ* is ≈ 10% smaller than the true Jaccard. As *g* increases, the bias decreases, e.g. for *g* = 100 and *L* =10, 016, *Ĵ* is 4% smaller than the true Jaccard.

## 4 Overview of Theorem 1 proof

Due to space constraints, we will focus only on the main theorem (Theorem 1) in the main text, providing the intuition and giving an overview of the technical highlights. The proofs of all the theorems, as well as all the building blocks, are deferred to the Appendix. Our main technical novelty is the derivation of a mathematical expression, 𝒞(*A, B*; *w*), that approximates the expected value of the size of the minimizer intersection *Î* (*A, B*; *w*) between two sequences *A* and *B*.

### Lemma 1.

𝒞(*A, B*; *w*) ⩽ 𝔼[*Î* (*A, B*; *w*)] ⩽ 𝒞(*A, B*; *w*q) + 2.

𝒞(*A, B*; *w*) is function of *w, L*, and of the *k*-mer-matching between *A* and *B*. In particular, when these parameters are known, then 𝒞(*A, B*; *w*) can be easily computed. We define 𝒞(*A, B*; *w*) formally in Appendix A.1, since it requires the introduction of additional notation. In Section 4.1, we give a high level proof of overview of Lemma 1 that does not require the definition of 𝒞.

To prove Theorem 1, we first use Lemma 1 to approximate the value of 𝔼 [*Ĵ*(*A, B*; *w*)].

### Lemma 2.

*Let w* ⩾ 2, *k* ⩾ 2, *and L* ⩾ 7(*w* + 1) *be integers. Let A and B be two duplicate-free sequences, each consisting of L k-mers. Then there exists* 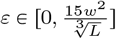 *such that*

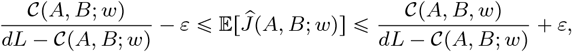

*where d* = 4/*w* +1.

Section 4.2 provides a sketch of the proof. Finally, to prove Theorem 1, we show that

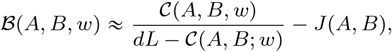

up an additive error that vanishes as the number of *k*-mers growths; when combined with Lemma 2 this approximation yields Theorem 1 immediately. In the following subsection, we will use *Î* as shorthand for *Î*(*A, B*; *w*); we will similarly use *Û,Ĵ, 𝒞*.

### 4.1 Lemma 1

In this section, we give an intuition for the proof of Lemma 1 and for where 𝒞(*A, B*; *w* comes from. Let 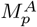 be the indicator random variable for the event that *A*_*p*_ is a minimizer in *A*. The expected size of the minimizer intersection can then be written in terms of 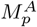 as follows:

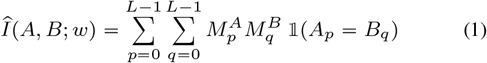

Here, we use 𝟙 in as an indicator function, i.e. 𝟙(*A*_*p*_ = *B*_*q*_) is 1 if *A*_*p*_ = *B*_*q*_ and 0 otherwise. Next, we use the notion of a charged window from [34, 20]. Given a position *p* ∈(0, *L* − 1) we say that *p charges* an index *i* if *i* ∈[max{−1, *p* − *w*}, *p* − 1], *a*_*p*_ = min{*a*_*i*+ 1_, …, *a*_min(*L*−*w*−1,*i*+*w*_)} and either *i* = max{−1, *p*−*w*} or *a*_*i*_ < *a*_*p*_. Figure 2 illustrates the definition. For *p* ∈ [0, *L* − 1] and *i* ∈ [−1, *L* − *w* − 1] we define 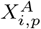 as an indicator random variable for the event that index *i* is charged by position *p*.

**Fig. 2:**
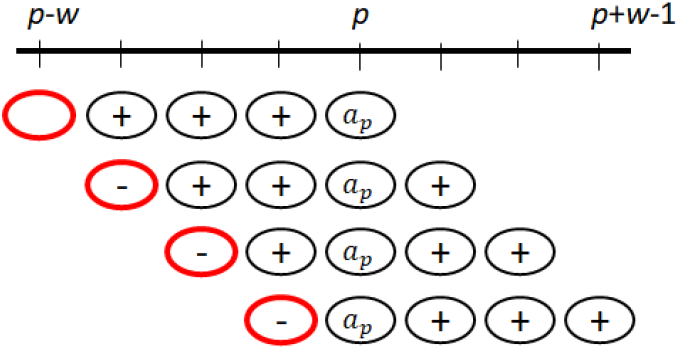
Illustration of charging. Each row shows a possible way that position *p* can charge an index, with *w=*4. A minus sign indicates the value is less than *a*_*p*_, a plus sign indicates the value is larger than *a*_*p*_, and no sign indicates that it does not matter. The circle at the index that is charged is shown in bold red. Note that no two rows are compatible with each other, i.e. every row pair contains a column with both a plus and a minus. As a result, the index that gets charged is unique.

The following fact was already shown in [34] and states that a minimizer charges exactly one window; Figure 2 shows the intuition behind it.

Fact 1. *Let p* ∈ [0, *L* − 1]. *Position p is a minimizer in A iff there exists a unique i* ∈ [−1, *L* − *w* − 1] *such that p charges index i. In other words*,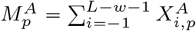.

Let us assume for the sake of simplicity and for this section only that *A* and *B* are padded. This allows us to combine Eq. 1 with Fact 1 while avoiding edge cases and get:

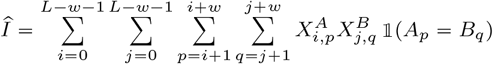

Applying linearity of expectation, the law of total probability, and the uniformity of the hash value distribution, we can show that

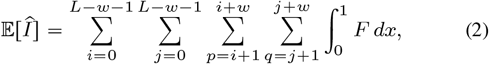

Where

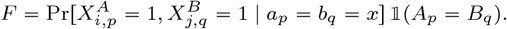

To derive the value of the probability term *F*, let us fix *p* and *q* such that *A*_*p*_ = *B*_*q*_ and fix *a*_*p*_ and *b*_*q*_ to be some value *x*. Observe that in order for 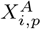 and 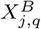 to both be one, there are certain positions that need to have a hash value less than *x* (which happens with probability *x* for each position) and certain positions that need to have a hash value more than *x* (which happens with probability 1 − *x* for each position). The hash values are pairwise independent, unless the two positions are in the *k*-mer-matching; in that case, the hash values are forced to be identical. If 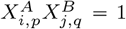 imply contradictory values for at least one position, then *F* is zero. Otherwise, let *α* be the number of hash values that need to be less than *x*, but counting matched pairs only once. Similarly, let *β* denote the number hash values that need to be more than *x*, counting the matched pairs only once. Then,

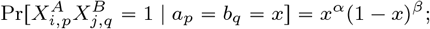

Figure 3 gives some examples.

**Fig. 3:**
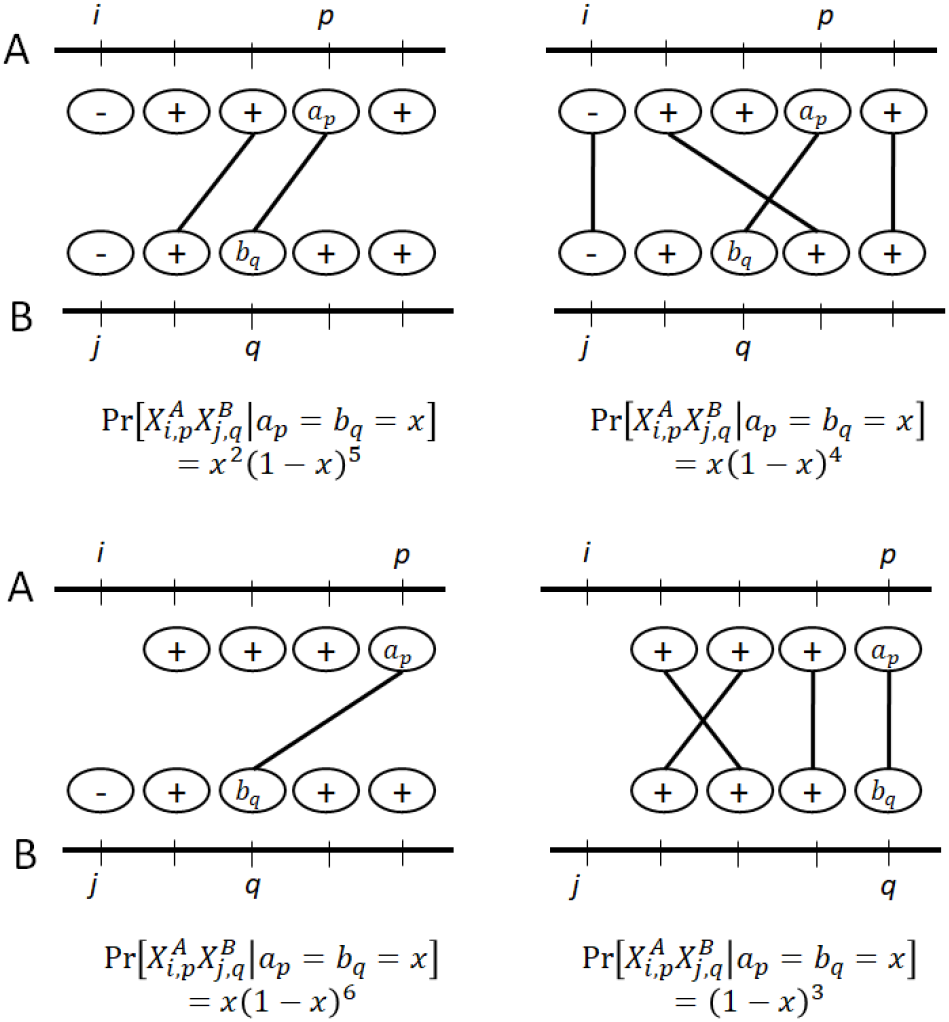
Some examples of 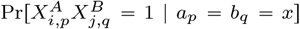, with *w* = 4. The two horizontal lines correspond to sequences *A* and *B*, and a circle corresponds to a *k*-mer whose value is relevant to the probability. The lines between *A* and *B* show the *k*-mer-matching, i.e. they indicate that the corresponding *k*-mers are the same. A plus or minus sign at a position reflects that the hash value must be greater or less than *x*, respectively.

Observe that 0 ⩽ *α* ⩽ 2 and 0 ⩽*β* ⩽ 2(*L* − 1). Therefore, the number of distinct terms in the summation of Eq. 2 is at most 6(*L* −1). The number of times each term is included in the summation is the number of *i, j, p, q* that induce the corresponding values of *α* and *β*. In Appendix A.1, we formalize this notion using *configuration counts*; but, for the purposes of intuition, it suffices to observe that Eq. 2 reduces to a function of the *k*-mer-matching, *w*, and *L*. We call this function 𝒞(*A, B*; *w*) and then obtain Lemma 1.

### 4.2 Lemma 2

In this section, we will prove Lemma 2, though we defer the proofs of the building blocks to the Appendix. Lemma 1 gives a tight approximation of 𝔼[*Î*] in terms of 𝒞. Now, we need to do the same for 𝔼[*Û*].

#### Lemma 3.

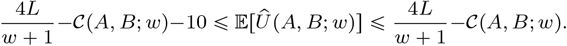

Now, with Lemmas 1 and 3, we can approximate 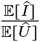. The next step is to show that this ratio of expectations is a good approximation for the expectation of the ratio 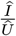, since 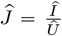. For this, we require asymptotically tight bounds on the variances of the random variables *Î*and *Û*

#### Lemma 4.

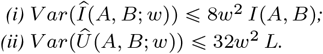

By isolating the central part of the distributions and bounding the effect of the tails using Chebyshev’s inequality [22], we then obtain the following approximation for 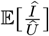.

#### Lemma 5.

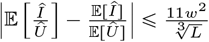.

We now have the components to prove Lemma 2.

Proof (Lemma 2). For the lower bound, we note that

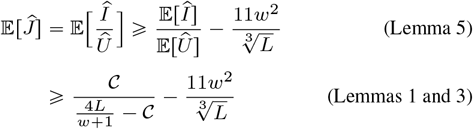

as claimed. For the upper bound, from Lemma 5, we know that

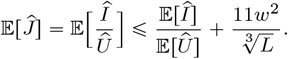

The bounds from Lemmas 1 and 3 imply

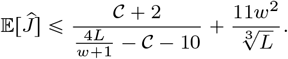

To complete the proof, we require two additional (and straightforward) bounds.

Fact 2.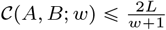.

Fact 3. *For all y* > 20 *and* 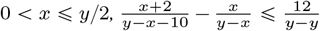

Letting *x* = 𝒞 and 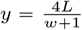, we have 0 < *x* ⩽*y*/2 and *y* > 20 (since *L* ⩾ 7(_*w*+1_) and so

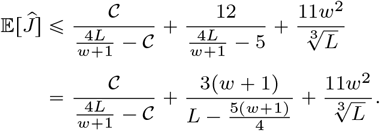

Plugging in *w* + 1 ⩽ *L*/7 and then using the fact that *w* ⩾ 2, we get

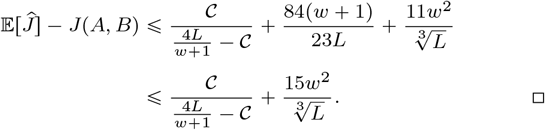

## 5 Empirical results

### 5.1 Experimental setup

We use two different models to generate sequence pairs. In the *unrelated pair* model, we take a desired Jaccard value *j*, set 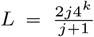, and independently and randomly generate two duplicate-free strings *A* and *B* with *L k*-mers. We chose *L* in this way so that under the assumption that *A* and *B* are uniformly chosen, *j* is the expected value of *J* (*A, B*), over the randomness of the generative process. While such string pairs are unlikely to occur in practice for higher values of *j*, they allow us to observe the bias of unrelated pairs for whole range of Jaccard similarities. In the *related pair* model, *A* is a randomly selected substring of *E. coli* [8] with *L k*-mers. String *B* is created by sweeping along *A*, at each position deciding with probability *r*_1_ whether to mutate and then choosing a new nucleotide from those that would not create a duplicate *k*-mer. More details about the handling of special cases are in Appendix A.6

For each model, we generated 50 *hash replicates* hash function (unless otherwise noted) where each replicate uses a different seed for the hash function. We then report 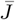, which is the average of *Ĵ* over the hash replicates and is the empirical equivalent of 𝔼[*Ĵ*]. We used the hash function that is part of minimap2 [16], since the idealized hash function we assumed for the convenience of our theoretical proofs is not practical in software. For the mutation model, we also generated some number of *mutation replicates*, where each replicate is the result of re-running the random mutation process. In any experiment, the same set of hash seeds were used for every mutation replicate. Scripts to reproduce our experiments are available on GitHub [26].

### 5.2 The extent of the empirical bias on real sequences

Figure 4 shows that there is considerable bias across a wide range of Jaccard values, for both related and unrelated sequence pairs. There are pairs of sequences with a dramatic bias, e.g. for unrelated pair with a Jaccard of 90%, the estimator gives only 44%. In more practically relevant cases, the bias can remain substantial; e.g. when the true Jaccard of related pairs is 76%, the estimator gives only 65% (when *w* = 200). The extent to which this bias is detrimental to the biological interpretation of the result depends on the downstream application. For example, using *Ĵ* to estimate the average nucleotide identity in order to build phylogenies, in the style of Mash [25], may be inadvisable.

**Fig. 4:**
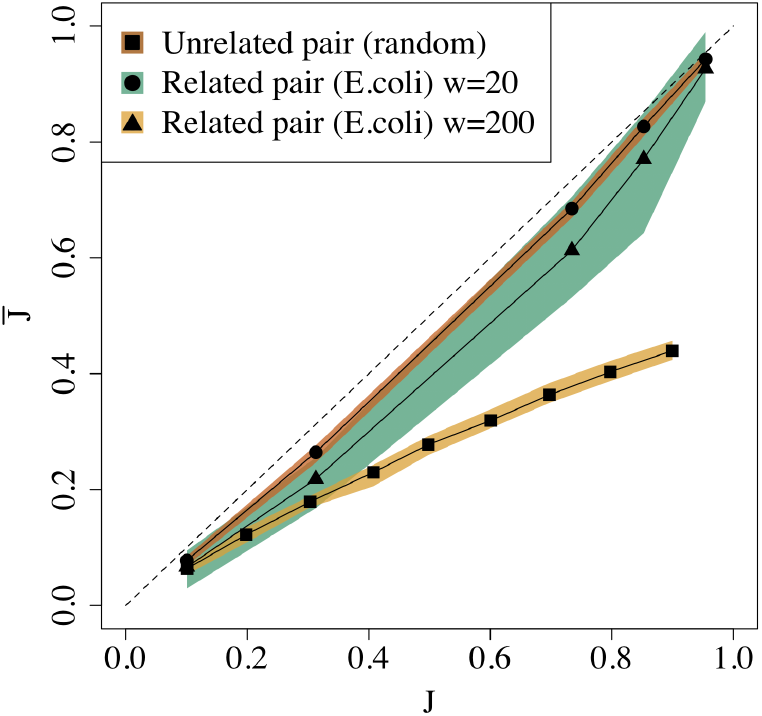
Empirical bias for unrelated and related sequence pairs. For the unrelated pairs, we used *w* 20 and *k =* 8 for *J* ⩾.4 and *k =* 7 for *J* ⩽..3. For related pairs, we set *k =* 16, *w* ∈{20, 200}, *L =* 10000, and *r*_1_ ∈{.001, .005, .01, .05, .1}, with one mutation replicate. The colored bands show the 2.5^th^ and the 97.5^th^ percentiles. The dashed line shows the expected behavior of an unbiased estimator, with 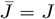.

Figures 4 and 5 show the extent to which the empirical bias depends on the window size *w*. Figure 4 shows that the bias for related pairs can be twice as large for *w* = 200 compared to *w* = 20. Figure 5 gives a more fine-grained picture and shows how the absolute bias for a related sequence pair increases with *w*. We note that it plateaus for larger values of *w*.

**Fig. 5:**
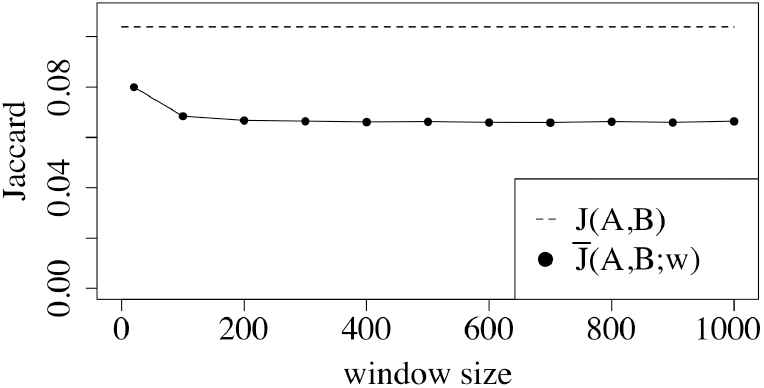
The effect of *w* on the empirical bias for a pair of related sequences as a function of the window size. Here, *r*_1_ = 0.1, *L =* 10, 000, *k =* 16, *w* ∈{20, 100, 200, …, 1000}, and there are 50 mutation replicates.

We also wanted to understand the extent of the bias in a scenario where the sequences are being compared as part of a read mapping process. To that end, we mimicked the behavior of the mashmap mapper [12, 13] by taking one arbitrary substring *A* from *E*.*coli*, with *L* = 1, 000, and comparing it to each substring *B* of *E*.*coli* with *L* = 1, 000. Figure 6 show that during the alignment process, we encounter the whole range of true Jaccard values, and, for each one, there is a substantial but not drastic bias in *JĴ*. Unlike the prediction of Theorem 2, the bias is sometimes positive; after further investigation, this happens because the *A* and *B* in this experiment are not always padded, which is a condition of Theorem 2.

**Fig. 6:**
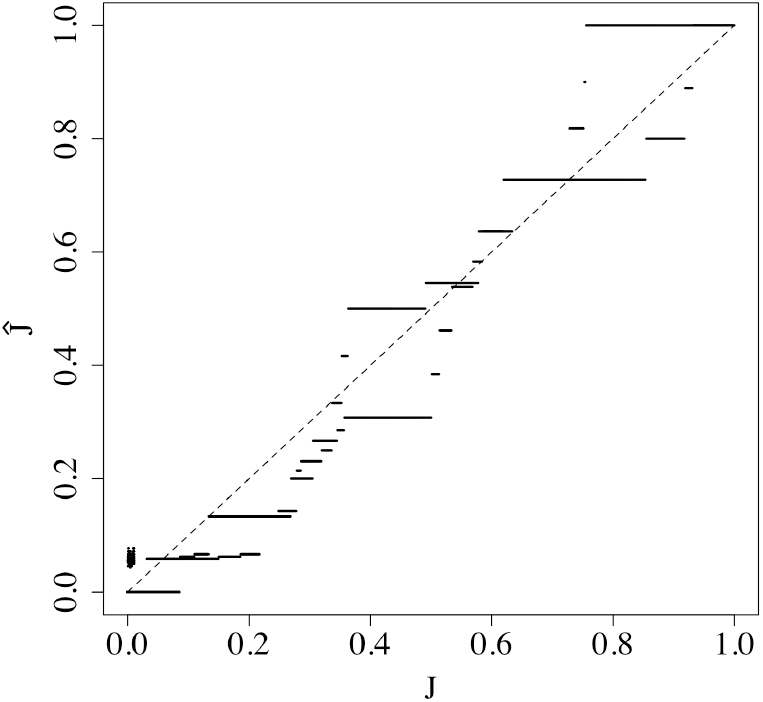
The empirical bias that occurs during a mapping process. Each point represents a comparison of a read *A* against a putative mapping location *B*. Note that the points visually blur into lines. We used *k =* 16 and window size *w =* 200 to match the default of mashmap. One hash replicate was used.

### 5.3 Effect of bias on mashmap sequence identity estimates

Mashmap is a read mapper that, for each mapped location, uses the Mash formula [25] to estimate the divergence (i.e. one minus the sequence identity) from *Ĵ*. It was previously reported that the Mash formula’s use of a Poisson approximation makes it inaccurate for higher divergence [33, 24], so before proceeding further, we modified mashmap to replace this approximation with the exact Binomial-based derivation (we derive the correction formula in Appendix A.6). We then simulated reads from *E*.*coli* with substitution errors to achieve a controlled divergence and mapped them back to the *E*.*coli* reference with mashmap (see Appendix A.6 for more details). We used *k* = 16 and mashmap automatically chose *w* = 200 as the window size.

Table 1 shows that even after our correction, the mashmap divergence had an error, e.g. for a true divergence of 5.00%, mashmap reported an average divergence of 5.71% – an error of 14%. To confirm that this remaining error was due to the minimizer sketch, we replaced the *Ĵ* estimator in mashmap with the true Jaccard. Table 1 shows that after this replacement, the remaining error was reduced by an order of magnitude, e.g. mashmap now reported an average divergence of 4.99%. We therefore conclude that the bias we observe in mashmap after the Binomial correction is dominated by the bias of *Ĵ*. In absolute terms, the *Ĵ* bias (about half a percentage point of divergence) may be acceptable for applications such as read alignment. However, for other applications (e.g. a fine grained analysis of sequence divergence), this bias may lead to downstream problems.

**Table 1.**
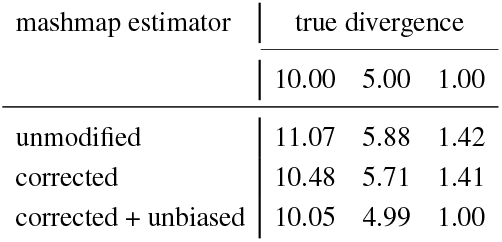
The median sequence divergence reported by mashmap, over 100 trials, for unmodified mashmap (first row), mashmap after Binomial-correction (second row) and, in addition, the removal of the *Ĵ* bias.

### 5.4 Empirical accuracy of our *ℬ* formula (Equation (3))

Theorem 1 predicts that our formula for *ℬ* (Equation (3)) approximates the empirical bias. To empirically evaluate the quality of this approximation, we measured the empirical error of Equation (3), which we define to be the absolute difference between the empirically observed bias 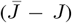 and ℬ. For the sequence pairs used in Figure 4, the empirical error is never more than 0.007 and roughly one to two orders of magnitude smaller than the bias itself (Tables 2 and 3). This held across three hash function families we tested: the one used by minimap2 [16], Murmurhash3 [23], and SplitMix64 [37]. Note that this robustness to different hash functions is not predicted by Theorem 1, which assumes an idealized version of a hash function which is collision free and maps uniformly to the real unit interval (in this case, none of the three functions map to the unit interval and Murmurhash3 is not collision free).

**Table 2.**
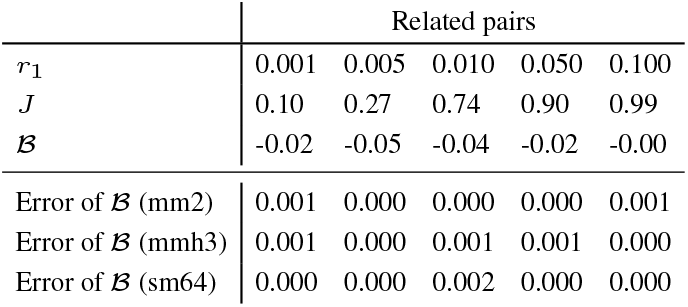
The empirical error of our theoretically predicted bias (Equation (3)) on the related pair sequences of Figure 4. The error is measured with respect to three different hash function families: the minimap2 hash function (mm2), the Murmurhash3 hash function (mmh3), and the SplitMix64 hash function (sm64).

**Table 3.**
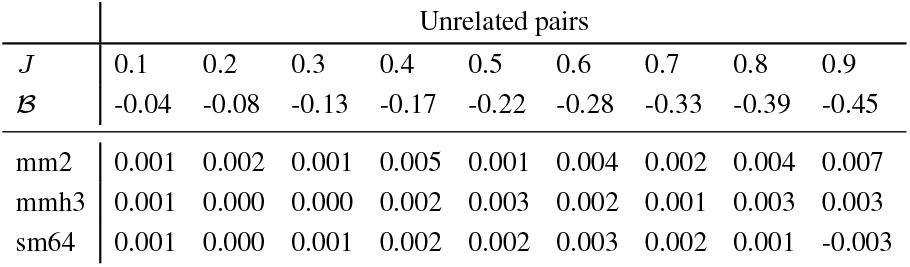
The empirical error of our theoretically predicted bias (Equation (3)) on the unrelated pair sequences of Figure 4. The error is measured with respect to three different hash function families: the minimap2 hash function (mm2), the Murmurhash3 hash function (mmh3), and the SplitMix64 hash function (sm64).

We measured the effect of increasing *w* and decreasing *L* on the empirical error for a related pair (Figure 7). The empirical error increases with *w* but remains almost two orders of magnitude smaller than the true Jaccard. For *L* ⩾ 1000, the empirical error is less than half a percent of the true Jaccard. Even for the smallest value of *L* (i.e. 100), the empirical error is only 2.6 percent of the true Jaccard. We conclude that Equation (3) is a high quality approximation for the empirically observed bias.

**Fig. 7:**
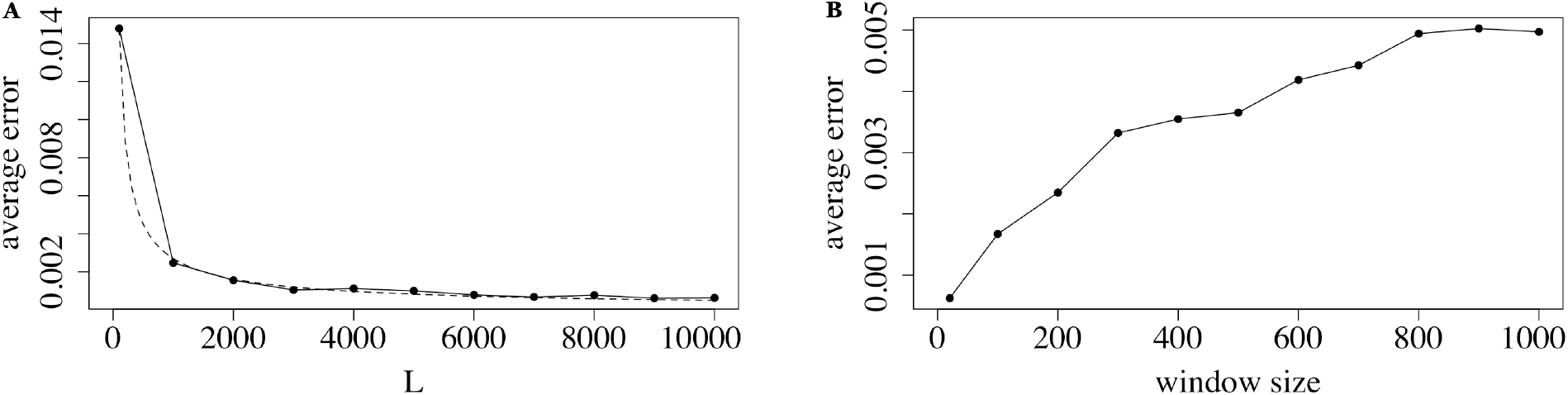
The effect of the window size *w* and sequence length on the empirical error of Equation (3). In panel A, we use the related pair model with 50 mutation replicates, *k =* 16, *w=* 20, *r*_1_ = 0.1, and *L* ∈ {100, 1000, 2000, …, 10000}. The y-axis shows the error of ℬ, averaged over the mutation replicates. The dashed line shows the best fit function of the form *αL*^*β*^, computed using the nls function in R. The average *J*, over the mutation replicates, is between .101 and .106, and the average empirical bias ranged between −0.023 and −0.027, depending on *L*. In panel B, we use the related pair model with 50 mutation replicates, *k* = 16, *L* = 10, 000, and *w* ∈ {20, 100, 200, …, 1000}. The average *J* is .104.

### 5.5 Accuracy of the *ε* bound to the approximation to Equation (3)

Theorem 1 states that the expected error of Equation (3) is at most 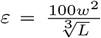. Since this is only an upper bound, we wanted to check the tightness with respect to *w* and to *L*. For *w* = 20 and non-astronomical values of *L, ε* > 1 and thus Theorem 1 gives no guarantee on the accuracy of the *B* term. Empirically, however, the error is small (Figure 7A), indicating that, at least for related pairs, *ε* is likely not a tight bound. To understand if the dependence on *L* is accurate, we found the best fit of a function of the form *αL*^*β*^ to the observed error curve in Figure 7A. The best fit was 0.44*L*^−0.74^, which indicates that our dependence on *L* in *ε* is not tight. One possible way to achieve this may be to use tighter concentration bounds than Chebyshev’s inequality inside the proof of Lemma 5 (leveraging the limited dependency between the events of *k*-mers being minimizers). Furthermore, Figure 7B suggests that the true error may be sub-linear in *w*, while *ε* has a *w*^2^ dependence. Thus our empirical results indicate that *ε* could potentially be improved for related sequences, though it may still be tight in the worst-case.

## 6 Discussion

In this paper, we showed that the minimizer Jaccard estimator suffers from bias and inconsistency, using both theoretical and empirical approaches. The bias can be drastic in some fairly artificial cases (i.e. unrelated sequences with high Jaccard) but remains substantial even on more realistically related pairs of sequences. Our theoretical results indicate that the bias cannot be removed by decreasing the window size (except for the pathological case when *w* = 1, where effectively there is no sketching done). We showed how the bias manifests in the mashmap read mapper as error in the reported sequence divergence. A future direction would be to derive the expected value of the bias ℬ in the simple mutation model of [2]; if ℬ reduces to a function of *w* without depending on the *k*-mer layout, then it could potentially be used to correct the bias in mashmap. Even if that were not possible, one could still use the estimator provided that an experimental evaluation determines that the observed bias is tolerable for the downstream application. On the other hand, the bias problems can be sidestepped altogether by using a similar but unbiased sketch, e.g. the modulo sketch [34]. Finally, we note that while we focus on bias in this paper, it is not the only theoretical property of importance for sketching; for example, there has been much exploration of different hash functions [20, 18, 7, 40, 9, 11, 14, 30] to reduce the density and/or to select *k*-mers that have desirable properties such as conservation or spread [35].

Our results also relate to the minhash minimizer Jaccard estimator (*Ĵ*_minhash_) described by [12]. In this variant, the set of *k*-mers in a minimizer sketch is further reduced by taking the *s* smallest values (i.e. their minhash sketch); the Jaccard estimator is then computed between these reduced sets. If the minhash sketch is taken using a different hash function than was used for computing minimizers, then the classical result of [3] implies that 𝔼[*Ĵ* _minhash_] = 𝔼[*Ĵ*]. This estimator would therefore suffer from the same bias that we have shown in this paper. If, on the other hand, the same hash values are reused, then the result of [3] is not applicable, because it assumes that the hash values being selected are uniformly random; in our case, the hash values being selected in the minhash step have already “won the competition” of being smallest in their window. Though we did not explore the bias of this variant of *Ĵ*_minhash_, it would seem surprising if the minhash step somehow magically unbiased *Ĵ*.

## A Appendix

In this appendix, we will prove the main theorems of the paper as well as provide experimental details to aid reproducibility.

### A.1 Matching configurations and the definition of 𝒞(*A, B*; *w*) and ℬ(*A, B*; *w*)

In this section, we define the notion of matching configurations and then use them to define 𝒞(*A, B*; *w*) and ℬ(*A, B*; *w*). As discussed in Section 3, the bias of *Ĵ* depends on the layout of the shared *k*-mers along the sequence. It turns out that the aspects of their sharedness that contribute to the bias are captured by the amount and location of *k*-mers that are shared between windows {*A*_*i*_, …, *A*_*i*+*w*_} and {*B*_*j*_, …, *B*_*j*+*w*_}, for any *i* and *j*.

Let us define *S* (*i, j*, 𝓁) ≜|{ *A*_*i*_, …, *A*_*i*+𝓁−1_ *B*_*j*_, …, *B*_*j+*𝓁−1_}|, i.e. the number of shared *k*-mers in the windows of length 𝓁 starting at positions *i* and *j* in *A* and *B*, respectively. We then define a *matching configuration* as a 5-tuple, written as

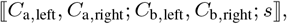

where *s* ∈ {0, …, *w*} and *C*_a,left_, *C*_a,right_, *C*_b,left_, *C*_b,right_ ∈ {0, 1, 2}. We then say that an index pair (*i, j*) with *i, j* ∈ [0, *L*−*w*−1] has configuration ⟦*C*_a,left_, *C*_a,right_; *C*_b,left_, *C*_b,right_; *s*⟧ if the windows {*A*_*i*+1_, …, *A*_*i*+*w*_} and {*B*_*j*+1_, …, *B*_*j*+*w*_} share *s k*-mers (i.e., *s* = *S*(*i* + 1, *j* + 1, *w*)) and

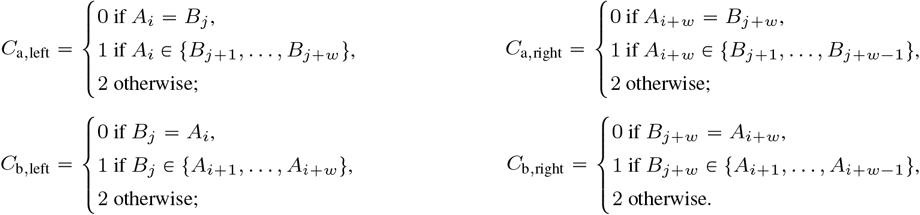

An index pair (*i, j*) has exactly one configuration, and not all configurations are possible; in particular, configurations where exactly one of *C*_a,left_ or *C*_b,left_ is zero, or exactly one of *C*_b,right_ and *C*_a,right_ is zero, are impossible. Figure S1 shows some examples of configurations. We may label configuration elements as sets (e.g. *C*_a,left_ = {0, 2}) to indicate all the configurations that can be formed using values from that set, except for impossible configurations. We use * as shorthand for the set {0, 1, 2} of all possible values. For example, ⟦*, 0; *, 0; *s*⟧ refers to the configurations ⟦0, 0; 0, 0; *s*⟧, ⟦1, 0; 1, 0; *s*⟧, ⟦2, 0; 1, 0; *s*⟧, ⟦1, 0; 2, 0; *s*⟧, ⟦2, 0; 2, 0; *s*⟧. For a configuration *C* we use *N* (*C*) to denote the number of pairs (*i, j*) such that the configuration of (*i, j*) is *C*.

In order to define ℬ(*A, B*; *w*), we define first the quantity 𝒞(*A, B*; *w*). Let 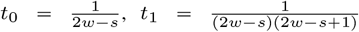, and 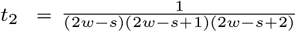.

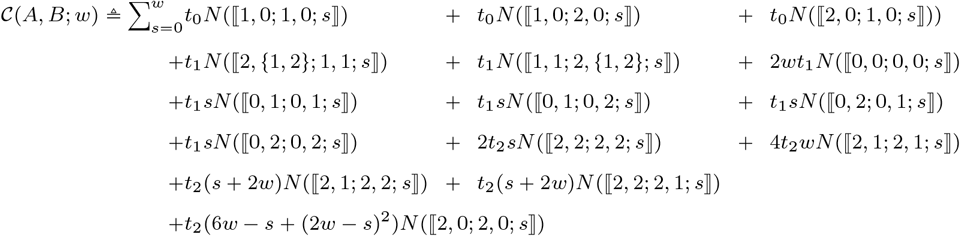

In particular, 𝒞 (*A, B*; *w*) is a linear combination of configuration counts, where each count is weighted by some function of its *s* value and *w*. We also define 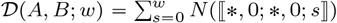 The term ℬ (*A, B*; *w*), which essentially determines the bias of the Jaccard estimator (see Theorem 1), is defined as follows:

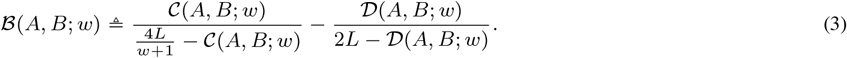

**Fig. S1:**
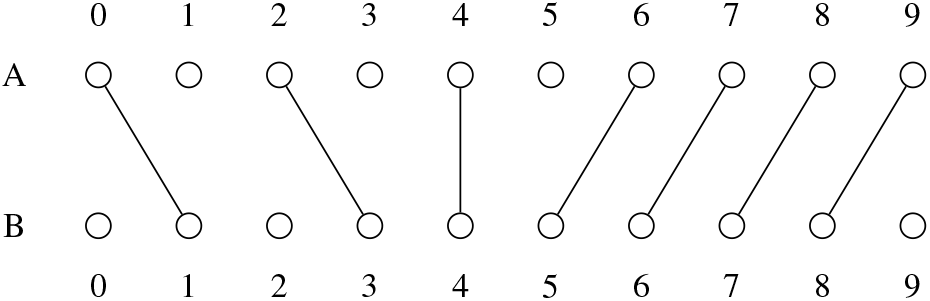
Configuration examples with *w* = 2: the pair (0, 1) has configuration ⟦0, 0; 0, 0; 1 ⟧; pair (4, 4) has ⟦0, 1; 0, 2; 1 ⟧; pair (7, 6) has ⟦0, 0; 0, 0; 2 ⟧.

### A.2 Proof of Theorem 1

In all the following, we will assume that *L* ⩾ 7(*w* + 1).

#### A.2.1 Approximating the minimizer union and intersection (Lemmas 1 and 3)

In this section, we will prove Lemmas 1 and 3. First, we recapitulate the proof of Fact 1 in our notation:

Fact 1. *Let* (∈ [0, *L* − 1]. *Position p is a minimizer in A iff there exists a unique i* ∈[−1, *L* −*w* −1] *such that p charges index i. In other words*,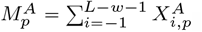.

Proof. Figure 2 gives the intuition for the proof. For the only if direction, suppose that *p* charges index *i*. Then, by definition of charging, *a*_*p*_ = min{*a*_*i*+1_, …, *a*_*i*+*w*_}, and so *p* is a minimizer. For the if direction, suppose that *p* is a minimizer in *A*. Consider the leftmost window in which it is a minimizer, i.e. the smallest *i*^′^ ∈ [*p* −*w* +1, *p*] such that 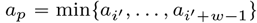. Since *i*^′^ is smallest, then either *i*^′^ = *p* −*w* +1 or 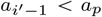. This is the definition of *p* charging index *i*^′^ − 1. For uniqueness, consider all the possible windows that *p* can charge, shown in Figure 2. They are all pairwise incompatible, i.e. there is at least one position that is simultaneously required to be larger than *a*_*p*_ and smaller than *a*_*p*_.

The expected value of 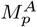 is called the *density* of the minimizer scheme, and we compute it exactly in the following Fact. We note that similar derivations of the density also appeared in [34, 28], but our proof accounts also for the edge cases.

Fact 4. *For p* ∈ [0, *L* − 1], *we have* 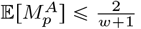. *More precisely*,

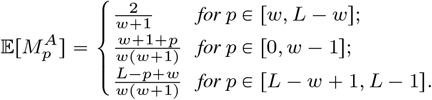

Proof. Let 𝓁 = max(−1, *p* − *w*) and *u* = min(*L* − *w* − 1, *p* − 1]. For *i* ∈[𝓁 + 1, *u*], we have 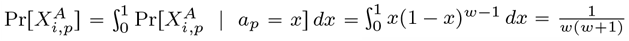. For *i=* 𝓁 we have 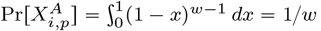.

By Fact 1, 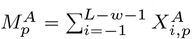.When *p* ∈ [0,*w* − 1], we have

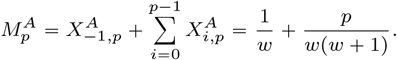

When *p* ∈ [0, *w* − 1], we have

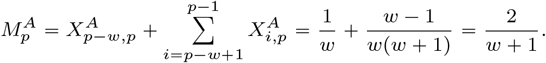

□

When *p* ∈ [*L*−*w* +1,*L* −1], we have

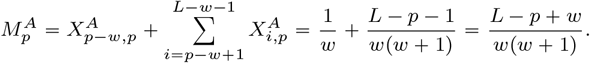

We are now ready to prove Lemma 1.

##### Lemma 1.

𝒞(*A, B*; *w*) ⩽ 𝔼[*Î*(*A, B*; *w*)] ⩽ 𝒞(*A, B*; *w*) + 2.

Proof. From the definition of *Î*(*A, B*; *w*) and Fact 1, we have

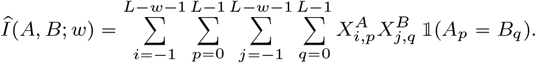

Observe that by definition of charging, 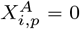 when *p* ∉ [*i* + 1, *i* + *w*]. Therefore,

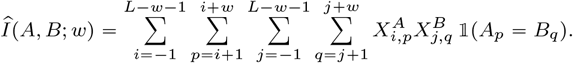

We can ignore some of the boundary terms associated with position −1 being charged without much loss in accuracy. Let

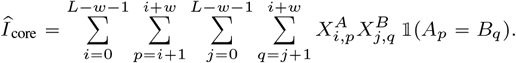

We claim that 𝔼[*Î*_core_] ⩽ 𝔼[*ÎA, B*; *w*) + 𝔼[*Î*_core_] + 2. The lower bound is immediate. For the upper bound, let us first separate out the terms of *Î* with *i* = −1 or *j* = −1:

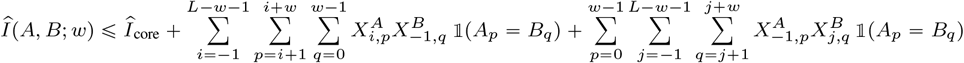

For the second term, observe that, by definition of charging, there is at most one value of *q* for which 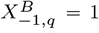. Then, since there are no repeated *k*-mers in *A* or *B*, there is at most one value of *p* for which *A*_*p*_ = *B*_*q*_. Finally, by definition of charging, there is at most one value of *i* for which 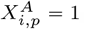. Hence the second term is at most one; by a symmetrical argument, the third term is at most one as well. This gives us the desired upper bound.

It now suffices to show that 𝔼[*Î*_core_] = 𝒞(*A, B*; *w*).

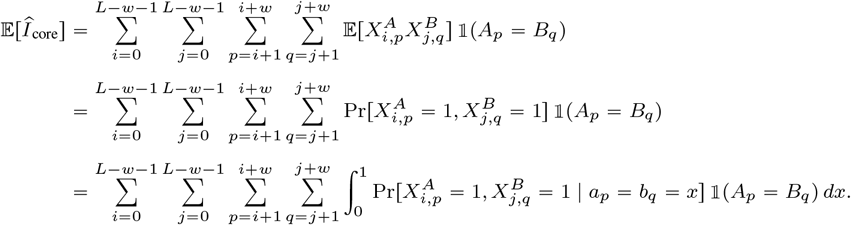

The probability 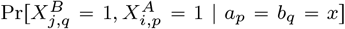 will depend on the configuration of the indices *i* and *j* and on whether *p* = *i* + *w* or *q* = *j* + *w*. Therefore, we rearrange the sums as follows. For a configuration *c*, we say that (*i, j*) → *c* when the indices *i* and *j* are in configuration *c*, so that

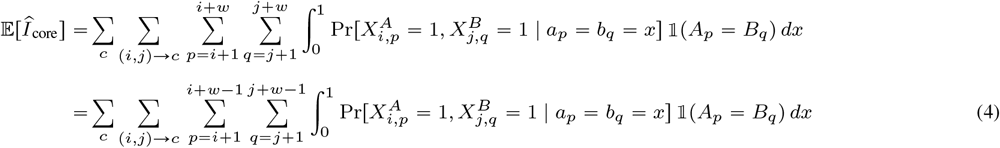

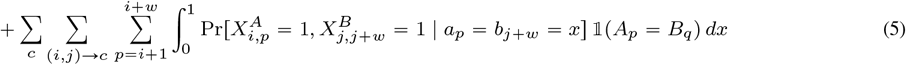

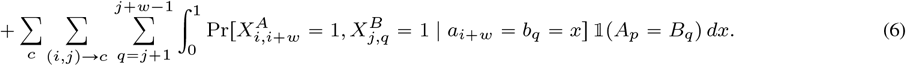

Figure 3 gives some examples to develop the intuition for what the inner term can evaluate to. We consider next each summation Equation (4), Equation (5), and Equation (6) separately. We start with Equation (5). Note that in this case the value of *q* is fixed to *j* + *w*, and so there is at most one value of *p* in the summation that is not 0 (since *A*_*p*_ = *B*_*q*_). We partition the space of all configurations into four possible cases: (i) *c* = ⟦+, 0; +, 0; *s*⟧, (ii) ⟦{0, 2}, *; *, 1; *s*⟧, (iii) *c* = ⟦1, *; *, 1; *s*], and (iv) *c* = ⟦*, *; *, 2; *s*⟧.

First note that for any *c*, we have 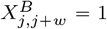 if and only if *b*_*j*+1_, …, *b*_*j*+*w*−1_ are each greater than *x*. In case (i) when *c* = ⟦*, 0; *, 0; *s*⟧, the only value of *p* for which the probability in Equation (5) is not zero is *p* = *i* + *w*. From the definition of charging, we have 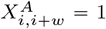 and 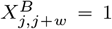 if and only if *a*_*i*+1_, …, *a*_*i*+*w*−1_, *b*_*j*+1_, …, *b*_*j*+*w*−1_ are each greater than *x*. The number of distinct *k*-mers in this sequence is 2*w* − 2 − *S*(*i* + 1, *j* + 1, *w* − 1) = 2*w* − 2 − *S*(*i* + 1, *j* + 1, *w*) + 1 = 2*w* − 1 − *s*. Therefore, 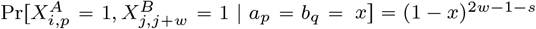 and

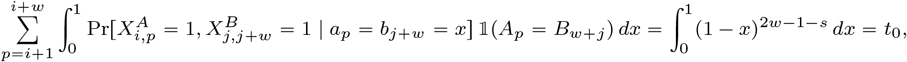

recalling that *t*_0_ 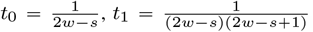, and 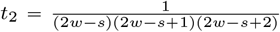. For case (ii) with *c* = ⟦{0, 2}, *; ., 1; *s*⟧, because *C*_b,right_ = 1, the only value of *p* for which the probability in Equation (5) is not zero belongs to [*i* + 1, *i* + *w* − 1]. From the definition of charging, we have 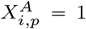 iff *a*_*i*_ < *x* and *a*_*i*+1_, …, *a*_*i*+*w*_, with the exception of *a*_*p*_, are all greater than *x*. As mentioned previously, we have that 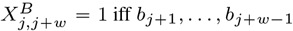 are each greater than *x*. Because *C*_a,left_≠ 1, we have *A*_*i*_ ∉ {*B*_*j*+1_, …, *B*_*j*+ *w*−1_). Therefore, we have one hash value (i.e. *a*_*i*_) that is less than *x*, and 2*w* − 2 − (*S*(*i* + 1, *j* + 1, *w*) −1) distinct hash values that are more than *x*. As a result,

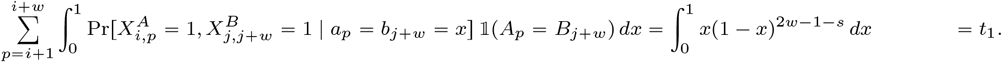

For next two cases (i.e., case (iii) and (iv)) we show that the sum is 0. When *c* = ⟦1, *; *, 1; *s*⟧, the fact that *C*_b,right_ = 1 means that *C*_a,right_≠ 0 which implies that *p* <*i* + *w* and that, if 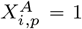, then *a*_*i*_ < *x*. The fact that *C*_a,left_ = 1 implies that *A*_*i*_ ∈ {*B*_*j*+1_, …, *B*_*j*+*w*_}. Therefore, one of the values of {*b*_*j*+1_, …, *b*_*j*+*w*_} is less than *x*, which makes it impossible that 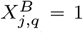. When *c* = ⟦*, * *, 2; *s*⟧, there is no value of *p* ∈ [*i* + 1, *i* + *w*} which satisfies *A*_*p*_ = *B*_*j*+*w*_, so 𝟙(*A*_*p*_ = *B*_*j*+*w*_) = 0. Putting all the four cases together, we have shown that the inner summation in Equation (5) is:

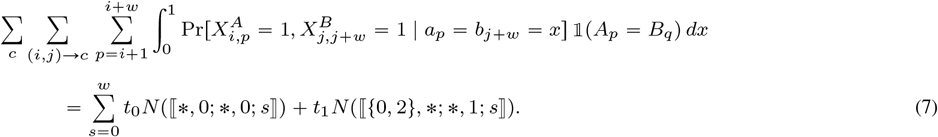

Deriving a closed form for Equation (6) is symmetric to Equation (5) with the exception that when *c* = ⟦ *, 0; *, 0; *s*⟧, there is no value of *q* in the range of the sum (i.e. *q* ∈ [*j* + 1, *j* + *w* − 1]) such that *A*_*i*+*w*_ = *B*_*q*_. Hence, for the inner summation in Equation (6), we obtain

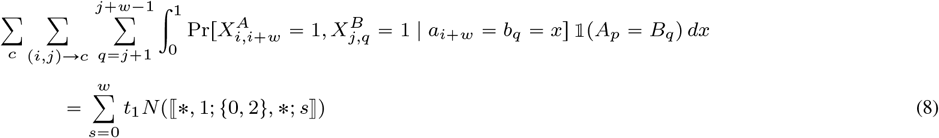

With a similar but more delicate case-by-case analysis, we also derive a closed form for Equation (4), whose proof we postpone until later.

Fact 5. *Let*

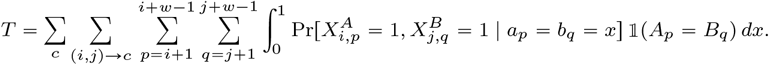

*Then*,

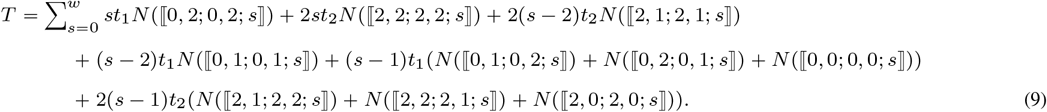

Finally, observe that summing Equation (7), Equation (8) and Equation (9) and then collecting the coefficients for each configuration, we obtain that *G* = 𝒞(*A, B*; *w*) as desired.□

We proceed with the proof of Fact 5.

Proof of Fact 5. For ease of notation, for a configuration *c* and a pair (*i, j*) → *c*, let

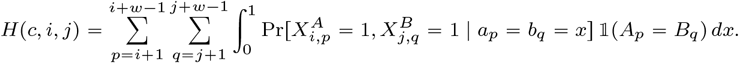

Since *p*≠ *i* + *w* and *q*≠ *j* * *w*, we have that 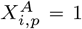 and 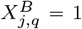 iff *a*_*i*_ < *x, b*_*j*_ < *x*, and *a*_*i*+1_, …, *a*_*i*+*w*_, *b*_*j*+1_, …, *b*_*j*+*w*_, with the exception of *a*_*p*_ and *b*_*q*_, are each greater than *x*. This corresponds to 2*w* − 1 − *s* hash values needing to be greater than *x*. What remains is to compute how many hash values need to be less than *x*.

We will partition the space of configurations into four possible cases: ⟦0, *; 0, *; *s*⟧, ⟦2, *; 2, *; *s*⟧, ⟦*, *; 1, *; *s*⟧, and ⟦1, *; *, *; *s*w. First, consider the case of *c* = ⟦0, *; 0, *; *s*⟧. In this case, *A*_*i*_ = *B*_*j*_. Therefore,

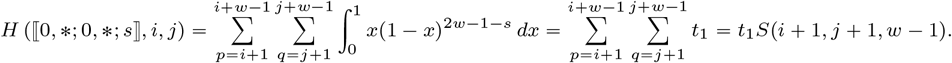

Next, consider the case of ⟦2, *; 2, *; *s*⟧. This case is exactly the same as *c* = ⟦0, *; 0, *; *s*⟧, except that *A*_*i*_ ≠ *B*_*j*_ and so

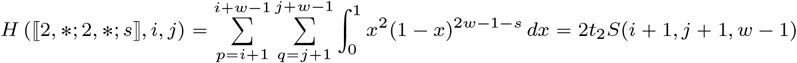

Next, observe that

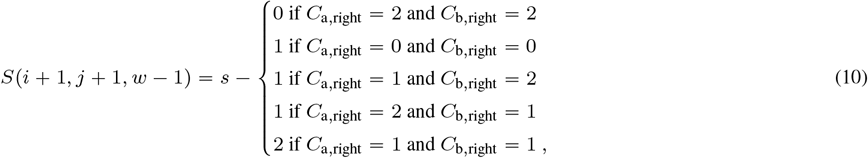

where recall that *s* = *S*(*i* + 1, *j* + 1, *w*). Therefore,

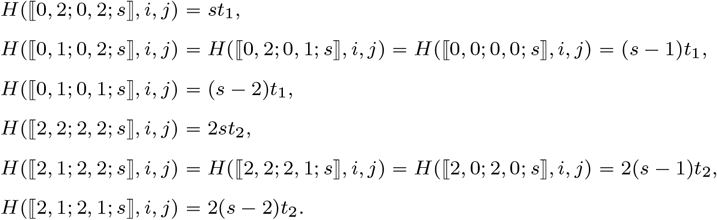

Now, when *c* = ⟦1, *; *, *; *s* ⟧, *A*_*i*_ ∈ {*B*_*j*+1_, …, *B*_*j*+*w*_}. However, we already argued that *a*_*i*_ > *x* and that *b*_*j*+1_, …, *b*_*j*+*w*_ are all at least *x*. Hence, we cannot have both 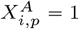 and 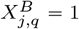, and this type of configuration does not contribute to the sum. The case of *c* = ⟦ *, *; 1, *; *s*⟧ is symmetric. Finally, observing that *T* = Σ_*c*_ *Σ* _(*i,j*)→*c*_ *H*(*c, i, j*), we combine all the cases to get the desired equality of the fact statement.

We now restate Lemma 3, whose proof is a direct consequence of Lemma 1.

##### Lemma 3

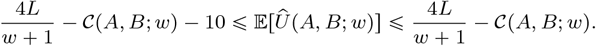

Proof. Recall that 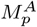 denotes the indicator random variable for *A*_*p*_ being a minimizer in *A*. Then

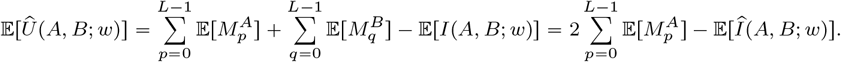

From Lemma 1, we know that 𝔼 [*Î*(*A, B*; *w*)] ⩾ 𝒞(*A, B*; *w*) and from Fact 4 we get that 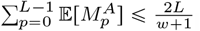. Combining these two facts, we deduce

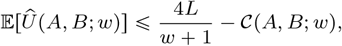

as desired. For the lower bound, from Fact 4 we can deduce that

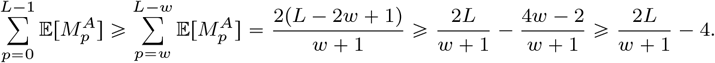

□

The lower bound then follows from Lemma 1.

#### A.2.2 Approximating the ratio of the minimizer union and intersection (Lemmas 4 and 5)

We begin this section with the proof of Lemma 4, where we obtain bounds for the variances of *Î*(*A, B*; *w*) and *Û* (*A, B*; *w*). Lemma 4.

*(i) Var*(*Î*(*A, B*; *w*)) ⩽8*w*^2^ *I*(*A, B*);

*(ii) Var*(*Û* (*A, B*; *w*)) ⩽ 32*w*^*2*^ *L*.

Proof. For ease of notation, we let *I* = *Î*(*A, B*) and *U* = *U* (*A, B*). If *p* is a position in *A*, then define *w*_*p*_ = {*A*_max{0,*p*−*w*+1}_, …, *A*_min{*p*+*w*−1,*L*−1}_} and, if *x* = *A*_*p*_, we say that the *k*-mers in *w*_*p*_ are *nearby x* in *A*.

We begin with part {*i*}. For ease of notation set *Î* = *Î*(*A, B*; *w*) and recall that

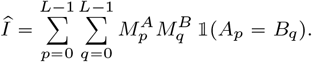

Then,

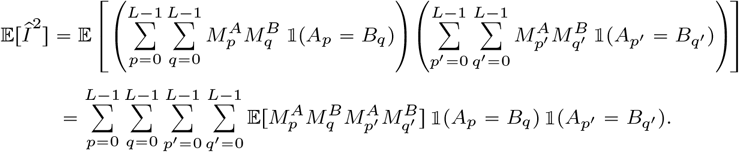

Observe that 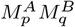and 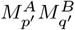 are independent if |*p − p*^*′*^| > 2(*w−*1),w_p_ ⋂ w_q_ ^′^*= Ø* and w_p_ ^′^ ⋂ w_q_ *= Ø*, since these four conditions guarantee that the two windows of size 2*w −*1 centered at *p* and *q* (which determine 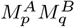) do not share *k*-mers with the two windows centered of size 2*w−*1 at *p*^*′*^ and *q*^*′*^ (which determine 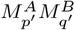).

Let *D* bet the set of tuples (*p, q, p* ^*′*^, *q*^*′*^) such that *p, q, p* ^*′*^, *q*^*′*^ ∈ r[0, *L*], *A*_*p*_ = *B*_*q*_, *Ap*^*′*^ = *Bq*^*′*^ and at least one of the following conditions hold: (i) |*p − p*^*′*^| ≤ 2(*w −* 1), (ii) |*q − q*^*′*^| ≤ 2(*w −* 1), (iii) *w*_*p*_ *⋂ wq*^*′*^ ≠ Ø, or (iv) *w*_*p*_^*′*^ *⋂ wq* ≠ Ø. That is, *D* contains all tuples (*p, q, p*^*′*^, *q*^*′*^) for which 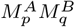 and 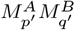 could be dependent, so that

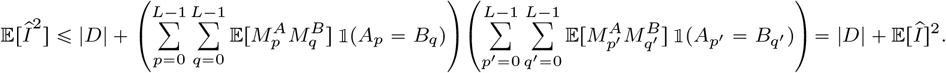

Then, *Var*(Î) = 𝔼 [Î]^2^ − 𝔼 [Î]^2^ ≤ |*D*| and it thus suffices to derive an upper bound for |*D*|. To do so, we will count the number of tuples that satisfy each of the conditions on the definition of *D* and add them together together to get an upper bound on |*D*|. For condition (*i*), there are *I* values of (*p, q*) such that *A*_*p*_ = *B*_*q*_, and for each one, there are 4*w* − 3 possible values of *p*^*′*^ such that |*p* − *p*^*′*^ | ≤ 2(*w* − 1). Then, for a given value of *p*^*′*^, there is at most one value of *q*^*′*^ that would satisfy *A*_*p*_^*′*^ = *B*_*q*_^*′*^. Therefore there are at most (4*w* − 3)*I* values of (*p, q, p*^*′*^, *q*^*′*^) that satisfy condition (i), i.e. *A*_*p*_ = *B*_*q*_, *A*_*p*_^*′*^ = *B*_*q*_^*′*^ and |*p* − *p*^*′*^ | ≤ 2(*w* − 1). By the same logic, there are at most (4*w* − 3)*I* values of (*p, q, p*^*′*^, *q*^*′*^) that satisfy condition (ii), i.e. *A*_*p*_ = *B*_*q*_, *A*_*p*_^*′*^ = *B*_*q*_^*′*^ and |*q* − *q*^*′*^ | ≤ 2(*w* − 1).

For condition (iii), again there are *I* values of (*p, q*) such that *A*_*p*_ = *B*_*q*_. Then, each *k*-mer *x ∈ w*_*p*_ can occur at most once in *B*, hence there are at most 2*w* − 1 values of *q*^*′*^ such that *x ∈ w*_*q*_^*′*^. Since |*w*_*p*_ | = 2*w* − 1, there are at most (2*w* − 1)^2^ values of *q*^*′*^ such that *w*_*p*_ *⋂ w*_*q*_^*′*^ ≠ Ø. For each value of *q*^*′*^, there is at most one value of *p*^*′*^ such that *B*_*q*_^*′*^ = *A*_*p*_^*′*^. Therefore, there are at most *I*(2*w* − 1)^2^ values of (*p, q, p*^*′*^, *q*^*′*^) that satisfy condition (iii), i.e. *A*_*p*_ = *B*_*q*_, *A*_*p*_^*′*^ = *B*_*q*_^*′*^ and *w*_*p*_ X *w*_*q*_^*′*^≠ Ø. By symmetric logic, the number of tuples that satisfy condition (iv) is also *I*(2*w* − 1)^2^.

Putting this all together, we get *V ar*(Î) ≤ |*D*| ≤ 2(4*w* − 3 +; (2*w* − 1)^2^)*I ≤* 8*w*^2^*I*, which completes the proof of part (*i*).

We prove part (*ii*) next. For a *k*-mer *x ∈ U*, let *U*_*x*_ be the indicator random variable for the event that *x ∈* Û (*A, B*; *w*). Let *D* be the set of all (*x, y*) pairs such that *x ∈ U, y ∈ U*, and *U*_*x*_ and *U*_*y*_ are dependent. Then,

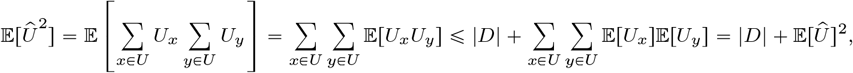

and *Var*(*Û*) = 𝔼 [Û ^2^] − 𝔼 [Û]^2^ ≤ |*D*|. It thus suffices to derive an upper bound for |*D*|. Let *x* and *y* belong to *U*. If *U*_*x*_ and *U*_*y*_ are dependent, then at least one of the following holds:

(i) One of the sequences (i.e. either *A* or *B*) contains both *x* and *y* at a distance of at most 2(*w* − 1).

*(ii) A* contains *x, B* contains *y*, and the nearby *k*-mers of *x* in *A* intersect with the nearby *k*-mers of *y* in *B*.

*(iii) B* contains *x, A* contains *y*, and the nearby *k*-mers of *x* in *B* intersect with the nearby *k*-mers of *y* in *A*.

We will count the possible number of (*x, y*) pairs that satisfy each of the conditions and use their sum as an upper bound on |*D*|. For (i), there are 2 choices for which sequence contains *x* and *y*, at most *L* choices for the position of *x*, and at most 4*w* − 3 choices for the position of *y*. Hence, there are at most 2*L*(4*w* − 3) choices for *x* and *y* that satisfy (i). For (ii), there are at most *L* choices for the position of *x*. If *y* satisfies the condition, then there must exist a *k*-mer *z* which is nearby to *x* in *A* and also nearby to *y* in *B*. There are at most 4*w* − 3 choices for *z*, and, for each of those choices, there are at most 4*w* − 3 locations for *y*. Hence, there are at most *L*(4*w* − 3)^2^ choices for *x* and *y* that satisfy (ii). Case (iii) is symmetrical to case (ii). In total then, |*D*| d 2*L*(4*w* − 3) + 2*L(*4*w* − 3)2 ≤ 32*w*^2^*L*.

With these bounds for the variances of Î(*A, B*; *w*) and Û (*A, B*; *w*) we can now prove Lemma 5.

##### Lemma 5.

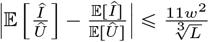

Proof. We start by introducing some convenient notation. Let 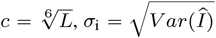 and.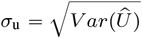 We say that Î and Û are *good* if their values lie in the range 𝔼[Î] ± *cσ*_i_ and 𝔼 [Û] ± *cσ*_u_, respectively; otherwise we say they are *bad*. Let 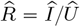. Note that 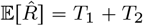, where

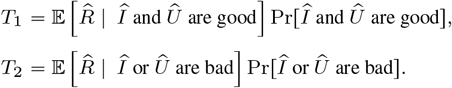

We will bound *T*_1_ and *T*_2_ separately. Observe that by Chebyshev’s inequality [22], the probability that Î is bad is at most *c*^−2^ and the same holds for Û. Hence, a union bound implies that Pr[Î or Û are bad] ≦ 2*c*^−2^. Since *I ≦ U*, 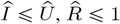, and we obtain the following bounds for *T*_2_:

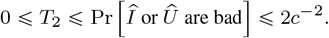

For *T*_1_, observe that

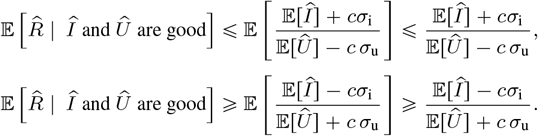

Also, since Pr[*Î* or *Û* are bad] ⩽ 2*c*^−2^, we have Pr[*Î* and *Û* are good] ⩾ 1 − 2*c*^−2^, and so

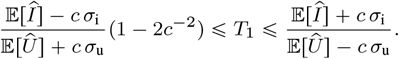

Now, observe that 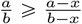, for 0 < *a* ⩽ *b* and 0 < *x* ⩽ *b* and 𝔼[*Î*] − *c σ*_i_ ⩽ 𝔼[*Û*] + *c σ*_u_, since 𝔼[*Î*] ⩽ 𝔼[*Û*] and *c* ⩾ 0.

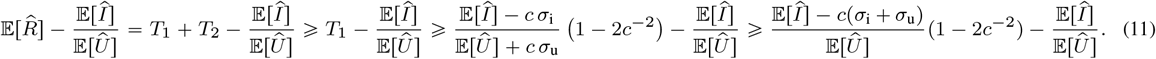

Observe that for all *x* > 0 and 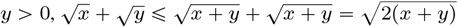. Then, using Lemma 4, we get:

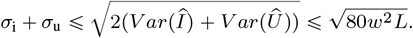

Furthermore, since every *w* consecutive *k*-mers have at least one minimizer, *Û* ⩾ *L*/*w*, and so

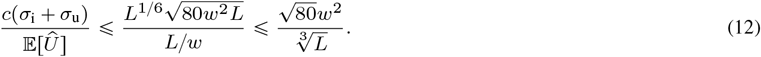

Plugging this bound into Equation (11) we get

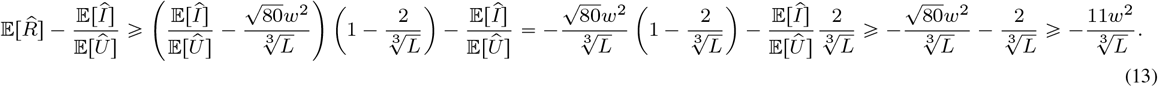

To derive the upper bound for 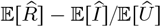, we first consider the case when 𝔼[*Û*] − 𝔼[*Î*] < *c* (*σ*_i_ + *σ*_u_). Under this assumption,

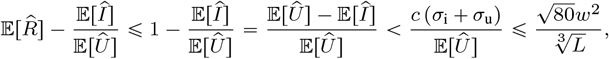

where the last inequality follows from Equation (12)

Now consider the case when 𝔼[*Û*] − 𝔼[*Î*] ⩾ *c* (*σ*_i_ + *σ*_u_). Using the fact that 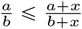, for 0 < *a* ⩽ *b* and *x* ⩾ 0, we obtain

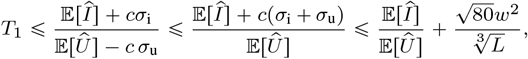

where the last inequality follows from Equation (12).

Putting the upper bounds on *T*_1_ and *T*_2_ together we get

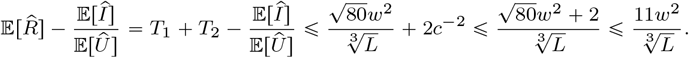

Combined with Equation (13) this implies the result. □

#### A.2.3 Proof of Theorem 1

To prove Theorem 1, we need to relate the bound on Ĵ(*A, B*; *w*) given by Lemma 2 to the values of *J* (*A, B*). We first express *J* (*A, B*) in terms of configuration numbers. Let 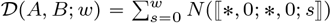. Note that, except near the start of the sequences, *A*_*i*_ = *B*_*j*_ if and only if (*i* − *w, j* − *w*) are in a configuration ⟦*, 0; *, 0; *s* ⟧. Therefore, 𝒟 [*A, B*; *w*] is approximately *I*(*A, B*). Formally, we can prove:

##### Lemma 6.

*If A and B are padded, then* 𝒟 (*A, B*; *w*) = *I*(*A, B*) *and* 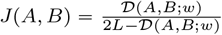 *More generally*,

i. 𝒟 (*A, B*; *w*) ⩽ *I*(*A, B*) ⩽ 𝒟 (*A, B*; *w*) + 2*w;*
ii. 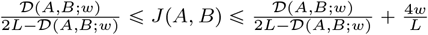

Proof. Observe that for *i* ∈ [*w, L* − 1] and *j* ∈ [*w, L* − 1], we have *A*_*i*_ *= B*_*j*_ if and only if (*i* −*w,j −w*) are in a configuration with *C*_a,right_ = *C*_b,right_ = 0. In the case that *A* and *B* are padded, then *I* = *𝒟* and 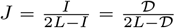. In general, the number of (*i,j*) pairs for which *A*_*i*_ *= B*_*j*_ and either *i* ∈ [0, *w* − 1] or *i* ∈ [0, *w* − 1] is at most 2*w*. Hence 𝒟 ⩽ *I* ⩽ 𝒟 + 2*w*. For the *J* lower bound, 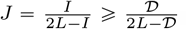. For the *J* upper bound, 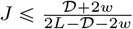. When 𝒟 + 2*w* ⩽ *L*, then

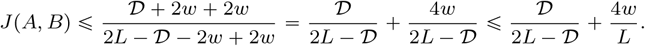

When *𝒟 +* 2*w > L*, then

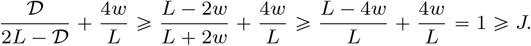

□

We note that it is possible to derive exact expressions for *I*(*A, B*; *w)* and *J*(*A,B*;*w*) for the non-padded case as well; however, doing so is not necessary for our purposes and would just introduce (even more) burdensome notation. Next, we need to prove two facts:

Fact 2. 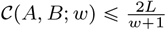

Proof. By Lemma 1, the definition of Î and Fact 4, we have 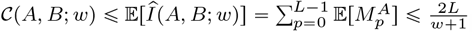. □

Fact 3. *For all y* > 20 *and* 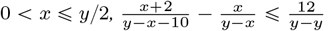.

Proof. Note that under the given assumptions, *y* − *x* ⩾ *y/*2 > 0 and *y* − *x* − 10 ⩾ *y*/2 − 10 > 0. Therefore,

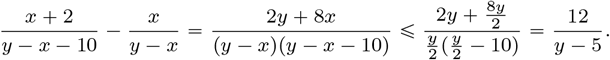

□

Now, we are ready to prove Theorem 1

##### Theorem 1.

*Let w ⩾* 2, *k* ⩾ 2, *and L* ⩾ 7 (*w +* 1*) be integers. Let A and B be two duplicate-free sequences, each consisting of L k-mers. Then there exists* 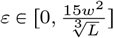 *such that*

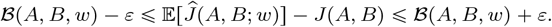

Proof. We prove the upper bound first. From Lemmas 2 and 6, we know that

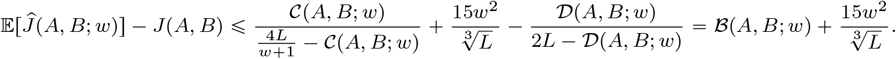

For the lower bound, we have

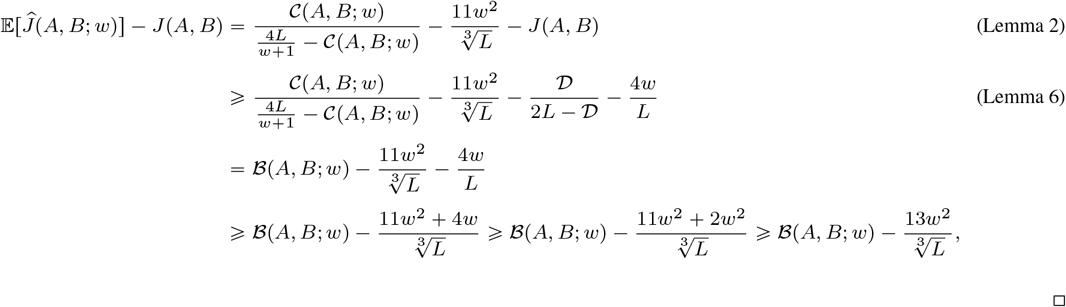

as claimed.

### A.3 Proof of Theorem 2

#### Theorem 2.

*Let w ⩾* 2, *k* ⩾ 2, *and L* ⩾ 7(*w+*1) *be integers. Let A and B be two duplicate-free padded sequences, each consisting of L k-mers. Then ℬ*(*A, B*; *w*) < 0 *unless J* (*A, B*) = 0, *we have ℬ*(*A, B*; *w*) = 0.

Proof. We omit the parameters *A, B* and *w* from the following for conciseness. Let 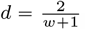. Observe that the following statements are equivalent:

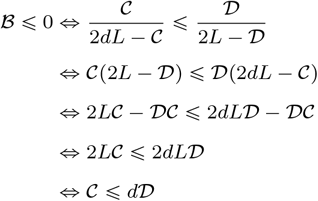

Note that for the second equivalence, we rely on the fact *ℬ*(*A, B*; *w*) is well defined and its denominators are not zero. In other words, 1) 2*L* − *D* > 0 because *𝒟* ⩽ *L* (by definition) and 2) 2*dL − C* > 0 because *C* ⩽ *dL* (by Fact 2).

We now need to show that *𝒞* ⩽ *d 𝒟*. We have

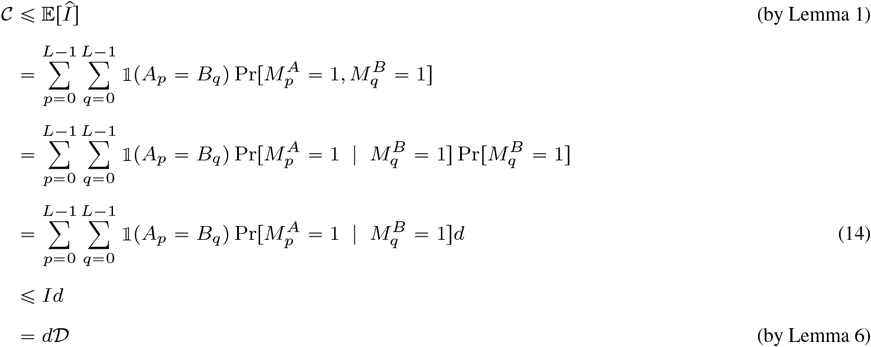

Note that Equation (14) follows because of the fact that *A* and *B* are padded and Fact 4. Next, observe that since all the terms in Equation (14) are positive, the only way to have equality with *Id* is if each term 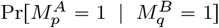 is 1. We claim this can only happen if there are no shared *k*-mers between *A* and *B*, i.e. when *J* (*A, B*) = 0. Otherwise, take the leftmost shared *k*-mer in *A*. The window to its left in *A* will be assigned hash values that are independent of the hash values in *B*; therefore, 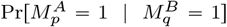 cannot be 1. Thus, if *A* and *B* share at least one *k*-mer, we get the stronger statement that 𝔼[*Î*(*A, B*; *w*)] < *Id*. This in turn implies that *𝒞* < *d 𝒟*, which propagates to imply that *ℬ* < 0.

### A.4 Proof of Theorem 3

#### Theorem 3.

*Let w ⩾* 2, *k* ⩾ 2, *and L* ⩾ 7(*w+*1) *be integers. Let A and B be two duplicate-free, padded, sparsely-matched sequences, each consisting of L k-mers. Then* 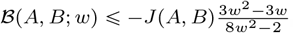.

Proof. This proof simply counts the configuration numbers and then applies definitions and Theorem 1. We will first count the configuration numbers. Let us call ⟬2, 2; 2, 2; ⟭ the *empty* configuration. Note that the terms involving the number of empty configurations cancel out in the equation for *𝒞* and hence we do not need to count them. Observe, by the condition of the theorem, that a configuration (*i, j*) that is non-empty must contain exactly one pair *p* ∈ [*i, i* + *w*] and *q ∈*[*j, j + w*] such that *A*_*p*_ *= B*_*q*_. Therefore, to count the number of non-empty configurations, it suffices to count, for every *p* ∈ [0,*L −*1] and *q* ∈ [0,*L* − 1] 1s such that *A*_*p*_ = *B*_*q*_, the types of configurations (*i, j*) for *i* ∈ [*p − w, p*] and *j* ∈ [*q − w, q*]. Following a case analysis, we get one configuration of ⟬2, 0; 2, 0; 1 ⟭, *w* − 1 configurations of ⟬2, 1; 2, 2; 1 ⟭, *w −* 1 configurations of ⟬2, 2; 2, 1; 1 ⟭, (*w −* 1)^2^ configurations of ⟬2, 2; 2, 2; 1⟭, one configuration of ⟬0, 2; 0, 2; 0 ⟭, *w* configurations of ⟬1, 2; 2, 2; 0 ⟭, and *w* configurations of ⟬2, 2; 1, 2; 0 ⟭. Recall that *I* = (*A, B*) is the number of shared *k*-mers between *A* and *B*. Summing over all *I* values of *p*, we then get the non-zero configuration number of non-empty configurations are

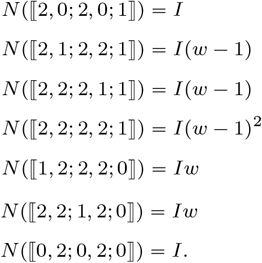

We then plug these into the definition of *𝒞* to get that *𝒞*(*A, B*; *w*) = *βI*, where 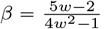. By Lemma 6, *𝒟*(*A, B*; *w*) = *I*. Let *d* ≜ 2/(*w*+1). Note that 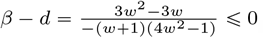. Using these facts, we can now derive

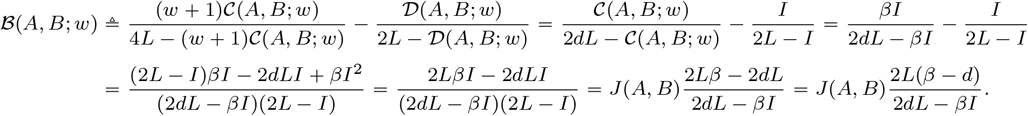

Note that because *β* − *d* ⩽0, *ℬ*(*A, B*; *w*) ⩽ 0. Then, using the fact that *β* > 0 and *I* > 0, we get

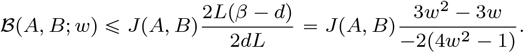

□

### A.5 Proof of Theorem 4

#### Theorem 4.

*Let* 2 ⩽ *w* < *k, g* > *w* + 2*k, and L* = *ℓg* + *k for some integer ℓ* ⩾ 1. *Let A and B be two duplicate-free sequences with L k-mers such that A and B are identical except that the nucleotides at positions k* − 1 + *ig, for i* = 0, …, *ℓ, are mutated. Then*,

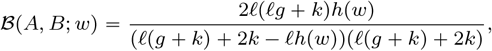

*where* 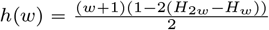 *and* 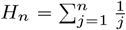 *denotes the n-th Harmonic number*. Proof. Let

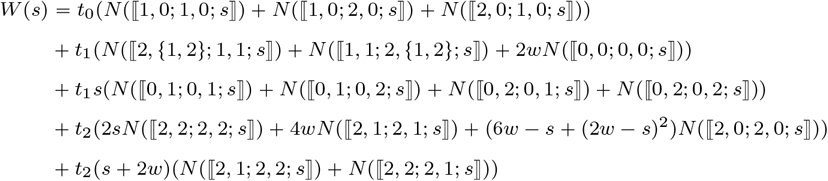

so that 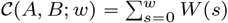. In our setting, the configuration counts are such that the following holds:

Fact 6.

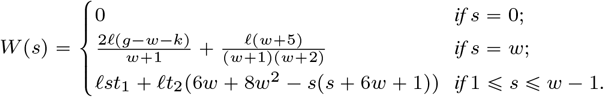

From this fact, which we prove later, we get that *𝒞*(*A, B*; *w*) = *dℓ*(*g* − *k*) + *ℓf* (*w*), where *d* = 2/(*w* + 1) and

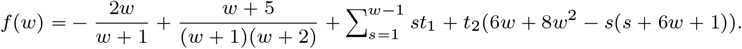

Note that since there are no matches in the first or the last *k*-mers and *k* ⩾ *w*, we have by Lemma 6 that *I* = |*A* ∩ *B*| = *𝒟*(*A, B*; *w*) = *ℓ*(*g* −*k*) and so

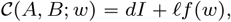

From the definition of *ℬ* (*A, B*; *w*), we then have

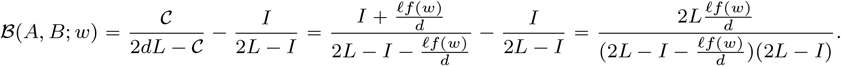

We also have the following closed form for *f* (*w*) (which we prove later).

Fact 7. *For n* ⩾ 1, *let*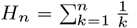. *Then, f* (*w*) = 1 − 2(*H*_2*w*_ − *H*_*w*_).

**Table S1.**
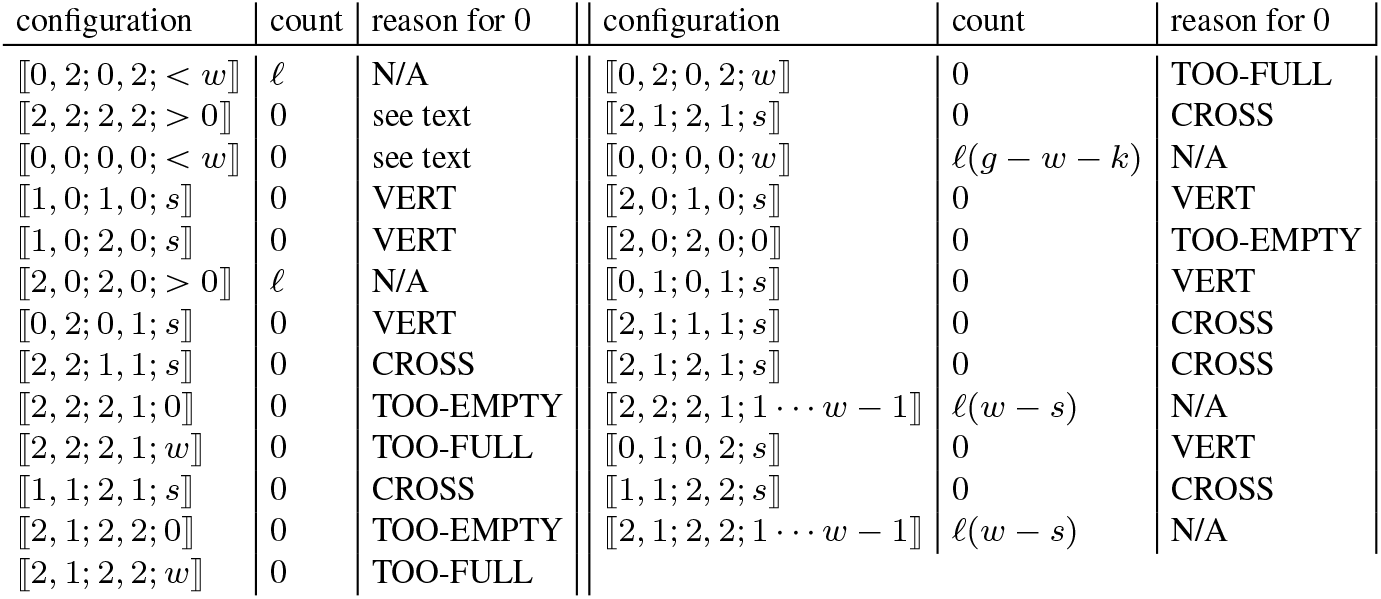
Non-empty configurations appearing in the definition of *𝒞*, along with their counts in the context of Theorem 4 as well as why the counts are zero, if applicable. The reasons are explained in the proof of Fact 8.

From this, combined with the facts that *L* = *ℓg* + *k* and *I* = *ℓ*(*g* − *k*), and letting 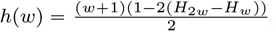, we get

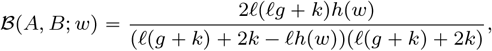

as claimed. □

It remains for use to provide the proofs of Facts 6 and 7. Fact 6 is a direct consequence of the following configuration counts.

Fact 8. *In the setting of* *Theorem* 4, *we have*

i. *N* (⟦0, 0; 0, 0; *w*⟧) = *l*(*g* − *w* − *k*);
ii. *N* (⟦0, 2; 0, 2; {0, …, *w* − 1}⟧) = *l;*
iii. *N* (⟦2, 0; 2, 0; {1, …, *w*}⟧) = *l;*
iv. *N* (⟦2, 1; 2, 2; {1, …, *w* − 1}⟧) = *l*(*w* − *s*);
v. *N* (⟦2, 2; 2, 1; {1, …, *w* − 1}⟧) = *l*(*w* − *s*).

*For any other configuration c that could contribute to 𝒞*(*A, B*; *w*), *we have N* (*c*) = 0 *or c* = ⟦2, 2; 2, 2; 0⟧.

Proof. We will refer to ⟦2, 2; 2, 2; 0⟧ as the *empty* configuration. Table S1 lists all non-empty configurations that appear in the definition of *𝒞*. Sometimes, a configuration type is further sub-divided according to different values of *s*. We will show that the counts in the table are correct, which will prove the Theorem.

The rows that whose reason is VERT have configurations that match ⟦*, *; 1, 0; *s*⟧, ⟦*, *; 0, 1; *s*⟧, ⟦1, 0; *, *; *s*⟧, or ⟦0, 1; *, *; *s*⟧. These configurations never occur because in our setting, all the matches are parallel to each other (i.e. if *A*_*i*_ = *B*_*j*_ and *A*_*i*′_ = *B*_*j*′_, then *j* − *i* = *j*′ − *i*′), while these configurations contain a 0 in one place (indicating that the matches are vertical, i.e. *A*_*i*_ = *B*_*j*_ implies *i* = *j*) and a 1 in another (indicated that the matching edges are angled, i.e. *A*_*i*_ = *B*_*j*_ implies *i* ≠ *j*). The rows whose reason is CROSS have a configuration that matches ⟦1, *; 1, *; *s*⟧, ⟦*, 1; *, 1; *s*⟧, ⟦1, 1; *, *; *s*⟧, or ⟦*, *; 1, 1; *s*⟧. These configurations never occur because the 1s indicate conflicting angles for the matches — they should either slant left (e.g. *i* > *j*) or right (e.g. *i* < *j*), but cannot do both. Note that for rows that could be categorized as both VERT and CROSS, the reason in the Table is arbitrarily chosen from those two. The rows whose reason is TOO-FULL have a configuration that matches ⟦*, 2; *, *; *w*⟧ or ⟦*, *; *, 2; *w*⟧. These configurations can never occur because the presence of the 2 indicates that either *A*_*i*+*w*_ or *B*_*j*+*w*_ is not involved in a match, making it impossible that *S*(*i* + 1, *j* + 1, *w*) = *w*. The rows whose reason is TOO-EMPTY have a configuration that matches ⟦*, *; *, {0, 1}; 0⟧ or ⟦*, {0, 1}; *, *; 0⟧. These configurations can never occur because the presence of the 0 or 1 indicates that either *A*_*i*+*w*_ or *B*_*j*+*w*_ is involved in a match, making it impossible that *S*(*i* + 1, *j* + 1, *w*) = 0.

By the definition of *A* and *B* from Theorem 4, we have alternating runs of *k* mismatches followed by *g* − *k* matches, with *k* mismatches at the end. Therefore, we have *ℓ* + 1 blocks of *k* mismatches, at *i* ∈ {*ig, …, ig* + *k* − 1|0 ⩽*i* ⩽*ℓ*u, and we have *ℓ* blocks of *g* − *k* matches, at *i* ∈ {*ig* + *k*, …, (*i* + 1)*g* − 1|0 ⩽*i* < *ℓ*}. We will refer to the latter as *match-blocks*.

Recall that configuration windows are of length *w* + 1. Because *k* > *w*, no window can contain matches from more than one match-block. Moreover, any configurations involving an *i* or *j* in the first match-block will occur again in each other match-block, at the same coordinates modulo *g*. Thus it is enough to consider only the first match-block, and multiply the resulting counts by *ℓ*. We therefore restrict ourselves to the first match-block in the following discussion, and note that the leftmost match is at position *k* and the rightmost match is at *g* − 1.

Let us consider the configurations that are ⟦2, 2; 2, 2; > 0⟧. In this case, *A*_*i*_ ≠ *B*_*j*_ and *A*_*i*+*w*_ ≠ *B*_*j*+*w*_, and there is some *i*′ ∈ [*i* +1, *i* +*w* −1] and *j*′ ∈ [*j* + 1, *j* + *w* − 1] such that *A*_*i′*_ = *B*_*j′*_. This match must be part of match block, and in our setting, a match block has width *g* − *k*. This is more than *w*, making it impossible that *A*_*i*_ ≠ *B*_*j*_ and *A*_*i*+*w*_ ≠ *B*_*j*+*w*_. Hence *N* (⟦2, 2; 2, 2; > 0⟧] = 0.

Let us consider the configurations that are ⟦0, 0; 0, 0; *s*⟧. In these configuration, *i* = *j, A*_*i*_ = *B*_*j*_, and *A*_*i*+*w*_ = *B*_*j*+*w*_. A configuration window of width *w* + 1 cannot span more than one match block, since *g* > *w*. Therefore, *A*_*i*+*δ*_ = *B*_*j*+*δ*_ for all 0 ⩽*δ* ⩽*w*. Hence, the number of configurations with *s* < *w* is 0. For *s* = *w*, Figure S2A shows all the configurations that are ⟦0, 0; 0, 0; *w*⟧. We have that *i* ∈ [*k, g* − *w* − 1], resulting in *g* − *w* − *k* possible windows with this configuration, in one match block

Let us consider the configurations that are ⟦0, 2; 0, 2; *s*⟧ for 0 ⩽ *s* ⩽ *w* − 1. In this situation, *A*_*i*_ = *B*_*j*_ and hence *i* = *j*. The match block containing this match ends before *A*_*i*+*w*_, since *A*_*i*+*w*_ ≠ *B*_*j*+*w*_ in this configuration. Then the rightmost match, *A*_*g*−1_ = *B*_*g*−1_, must be somewhere in the window, other than at *i* + *w*. To get *s* matches, *g* − 1 = *i* + *s* and thus *i* = *g* − *s* − 1. Therefore, *N* (⟦0, 2; 0, 2; *s*⟧) = 1 for each *s* ∈ [0, *w* − 1]. Figure S2B shows how this configuration looks like. The top and bottom drawings show the two end cases, while the middle drawing demonstrates the general case.

Let us consider the configurations that are ⟦2, 0; 2, 0; *s*⟧ for 1 ⩽ *s* ⩽ *w*. The case is mostly symmetric to the previous one. In this situation, *A*_*i*+*w*_ = *B*_*j*+*w*_ and hence *i* = *j*. The match block containing this match begins after *A*_*i*_, since *A*_*i*_ ≠ *B*_*j*_ in this configuration. The leftmost match in the match-block, *A*_*k*_, must be somewhere in the window other than at *A*_*i*_. To get *s* matches, *k* = (*i* + *w*) − (*s* − 1) and thus *i* = *k* − *w* + *s* − 1. Therefore *N* (⟦2, 0; 2, 0; *s*⟧) = 1 for each *s* ∈ [1, *w*]. Figure S2C shows how this configurations looks like. The top and bottom drawings show the two end cases, while the middle drawing demonstrates the general case.

Let us consider the configurations that are ⟦2, 1; 2, 2; *s*⟧ for 1 ⩽ *s* ⩽ *w* − 1. Figure S2D shows all the configurations. There are several possibilities for each *s*. For *s* = 3, the top and bottom drawings show the two end cases, while the middle drawing demonstrates the general case. Because *C*_a,right_ = 1, *A*_*i*+*w*_ ∈ {*B*_*j*+1_,…, *B*_*j*+*w*−1_} and *j* > *i*. Since *C*_a,left_ = *C*_b,left_ = 2, *A*_*i*_ ≠ *B*_*j*_, and the leftmost match in the match-block, *A*_*k*_, must be somewhere in the window, other than at *i*. To get *s* matches, *k* = (*i* + *w*) − (*s* − 1) and thus *i* = *k* − *w* + *s* − 1. The window for *B* can be positioned so that the leftmost match occurs in {*j* + 1, …, *j* + *w* − *s*}. Since this corresponds to *A*_*k*_, we have *k* ∈ {*j* + 1, *…, j* + *w* − *s*}, which can be restated as (*i* + *w*) − (*s* − 1) ∈ {*j* + 1,*…, j* + *w* − *s*}. We can in turn restate this as *i* ∈ {*j* − *w* + *s,…, j* − 1)} and thus *j* ∈ {*i* + 1,*…, i* + *w* − *s*)}. Therefore, *N* [⟦2, 1; 2, 2; *s*⟧) = *w* − *s* for each *s* ∈ [1, *w* − 1].

Finally, we consider the configurations that are ⟦2, 2; 1, 2; *s*⟧ for 1 ⩽ *s* ⩽ *w* − 1. This case is symmetrical to the above case, by swapping the roles of *A* and *B* in the definition of the configurations. Therefore, *N* (⟦2, 2; 1, 2; *s*⟧) = *w* − *s* for each 1 ⩽ *s* ⩽ *w* − 1. □

We are now ready to prove Fact 6.

Fact 6.

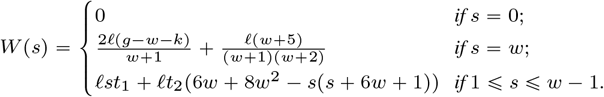

Proof. Let us consider first the *s* = 0 case. By Fact 8, the only two configurations with *s* = 0 and with non zero counts are ⟦2, 2; 2, 2; 0⟧ and ⟦0, 2; 0, 2; 0⟧. However, both of those terms are multiplied by *s* in *W* (0), hence we have *W* (0) = 0.

Let us consider next the *s* = *w* case. For this value of *s*, by Fact 8, we have *N* (⟦0, 0; 0, 0; *w*⟧) = *l*(*g* − *w* − *k*) and *N* (⟦2, 0; 2, 0; *w*⟧) = *l*; all other configurations that may contribute to *𝒞*(*A, B*; *w*) have zero counts.

At *s* = *w*, ⟦0, 0; 0, 0; *w*⟧ has coefficient 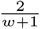 and ⟦2, 0; 2, 0; *w*⟧ has coefficient 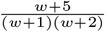. Hence

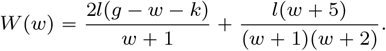

Finally, when 1 ⩽ *s* ⩽ *w* − 1, again by Fact 8, we have

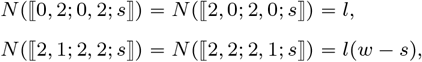

and all other configurations do not contribute to *W*. Now, the coefficient of *N* (⟦0, 2; 0, 2; *s*⟧) in *W* is *st*_1_, the coefficient of *N* (⟦2, 0; 2, 0; *s*⟧) in *W* is *t*_2_(6*w* − *s* + (2*w* − *s*)^2^), and the coefficient of *N* (⟦2, 1; 2, 2; *s*⟧) and *N* (⟦2, 2; 2, 1; *s*⟧) in *W* is *t*_2_(*s* + 2*w*). Combining this, we obtain

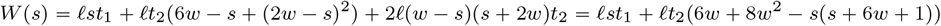

as claimed.

We conclude this section with the proof of Fact 7.

Fact 7. *For n* ⩾ 1, *let* 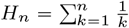. *Then, f* (*w*) = 1 − 2(*H*_2*w*_ − *H*_*w*_).

Proof. Recall that 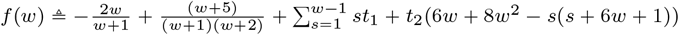. Let us rewrite *f* (*w*) as

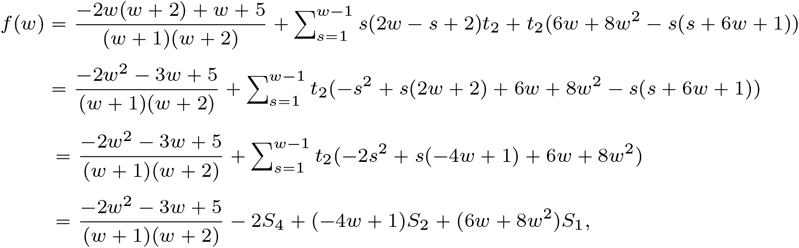

where 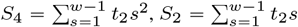 and 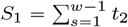.Let

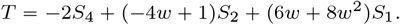

We will now reduce each of the sums.

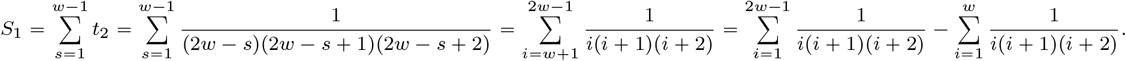

We now use the fact that 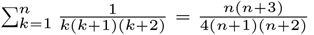, which can be derived via partial fraction decomposition or induction. Then,

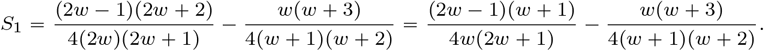

Proceeding similarly for the next component, we have:

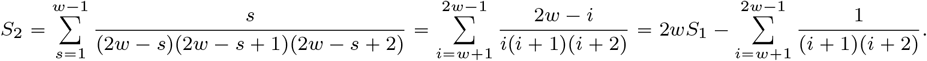

Recalling that 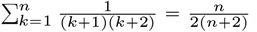, we get

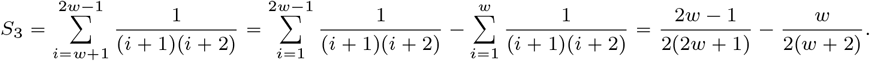

Hence

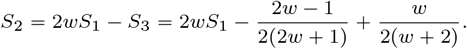

Finally,

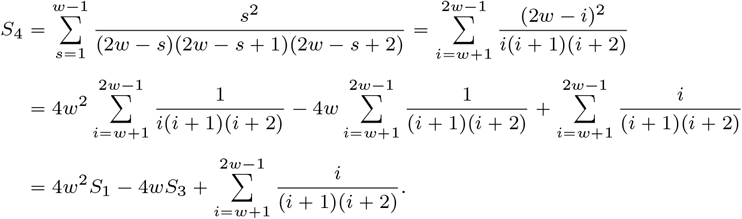

Using that 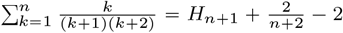 again via partial fraction decomposition or induction, we get

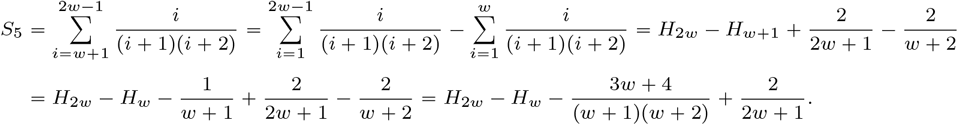

Thus,

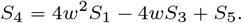

Combining all of this, we get

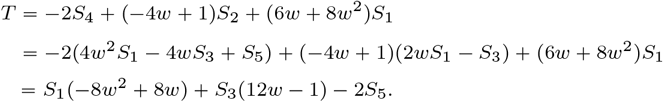

By using partial fraction decomposition, we can algebraically simplify each of the terms as follows:

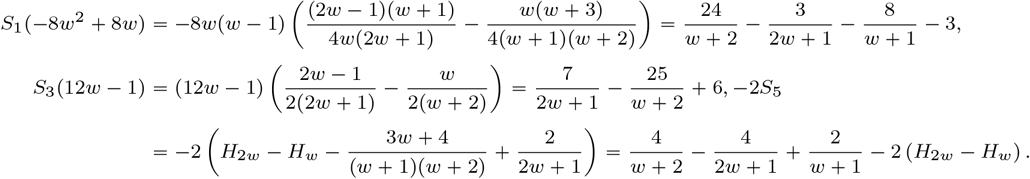

By plugging these expressions back into *T*, we get

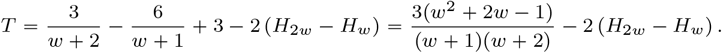

Now, we plug the value of *T* into *f* (*w*) and it finishes the proof,

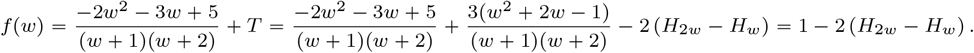

**Fig. S2:**
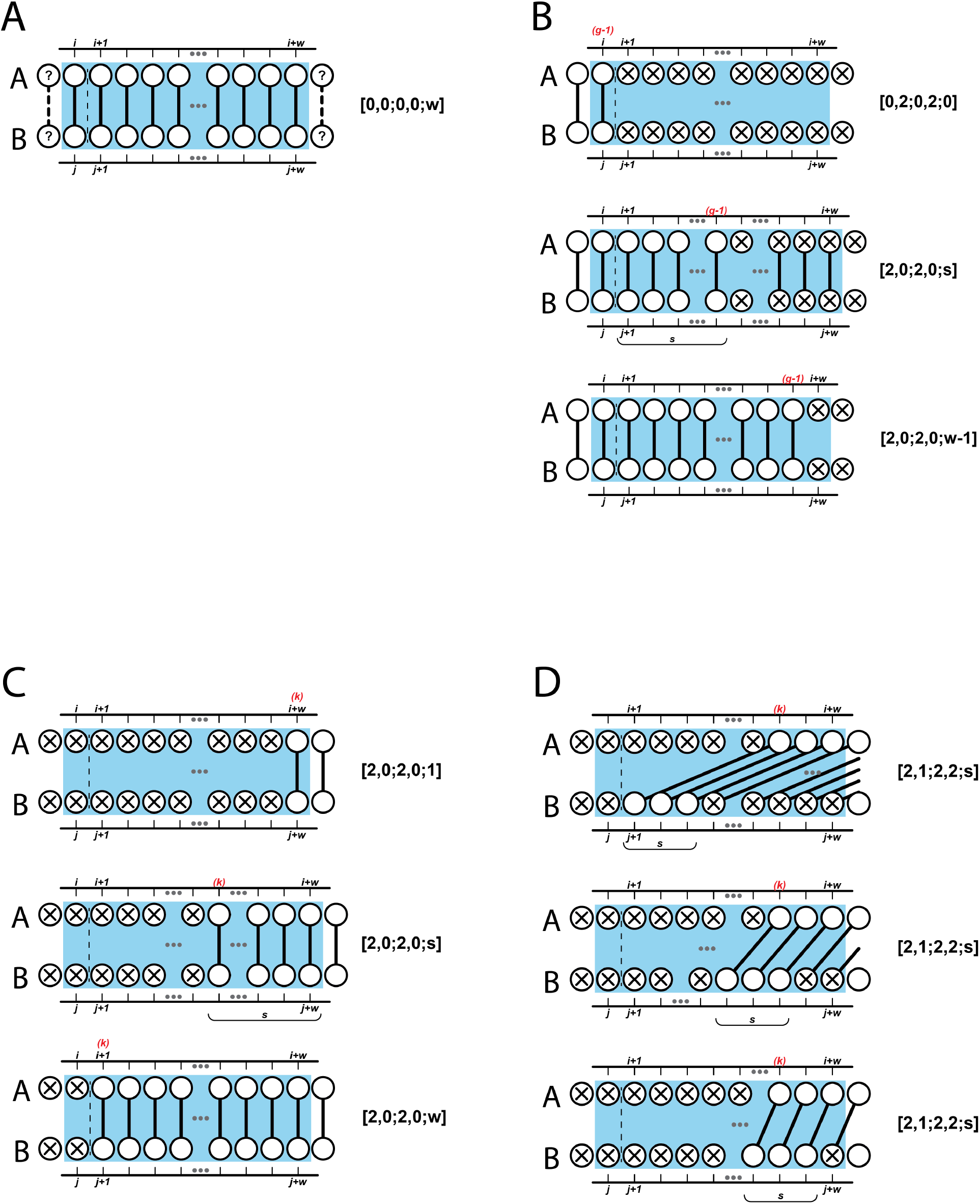
Some of the configurations with non-zero counts in Fact 8.

### A.6 Experimental details

In this section, we provide some experimental details to aid reproducibility. The scripts to reproduce our experiments are available on GitHub [26].

#### Generative models

When we generate an unrelated pair, we greedily extend each string from left to right. At each position, we choose, uniformly at random, one of the nucleotides that would not result in a *k*-mer we have already seen. If we get to a point where all the possible nucleotide extensions to a string are already present, we discard the string and start from the beginning. Though this sampling scheme is not guaranteed to terminate, we found that it always did in our experiments. We also verified that the Jaccard of the generated pair was close to the *j* that was used as a target. Under the assumptions that *A* and *B* are uniformly chosen, *j* should be the expected value under the generative process. Though it is not clear that the uniformity assumption holds in our generative process, we found that the true Jaccard was indeed very close to *j* in practice. In the related pair model, we also faced a possibility that after choosing to mutate a position, all the possible nucleotide substitutions would create a duplicate *k*-mer. In such a case, the position was left unchanged.

#### Mashmap divergence experiment

We sampled 100 substrings from the *E*.*coli* reference [8], each of length *L* = 10, 000 and, for each substring and for each *r*_1_ ∈ {0.90, 0.95, 0.99}, generated a “read” which was the substring with *r*_1_*L* positions randomly picked and mutated. We then mapped it with mashmap, and discarded any read for which mashmap did not correctly identify a unique and correct mapping location. Mashmap was run with default parameters of *k* = 16 and *w* = 200.

#### Correction formula to remove Poisson-approximation from Mash distance

Let *j* be the observed Jaccard. Let *A* and *B* be two sequences generated using a simple mutation process, i.e. a substitution is created at every nucleotide with a given probability *r*_1_ [2]. The method of moments [38] estimator for the sequence identity is *î* _mom_ = (1−*n*/*L*)^1/*k*^, where *n* is the observed number of mutated *k*-mers [2]. In the simple mutation model, the observed Jaccard *j* is related to *n* via 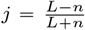, or, equivalently, 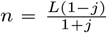 [2]. Putting this together, we get that 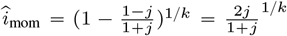. On the other hand, the Mash distance estimator is 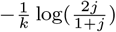 (Formula 1 in [12]), which equivalently translates to the identity estimator 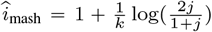. Combining the two, we get that 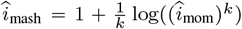. Solving for *î*_mom_, we get the final correction formula: 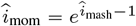.

#### Sliding read experiment

When choosing *A*, we avoided segments with any Ns or any duplicate *k*-mers. Any *k*-mers in *B* containing an N were hashed to the maximum hash value so as to avoid them being a minimizer. Also note that minimizers were computed separately for each *B*; thus, it is possible that the same *k*-mer might be a minimizer in one *B* but not a minimizer in a nearby *B*.

